# Tracking small animals in complex landscapes: a comparison of localisation workflows for automated radio telemetry systems

**DOI:** 10.1101/2024.04.14.589351

**Authors:** Cristina Rueda-Uribe, Alyssa J. Sargent, María Ángela Echeverry-Galvis, Pedro A. Camargo-Martínez, Isabella Capellini, Lesley T. Lancaster, Alejandro Rico-Guevara, Justin M. J. Travis

## Abstract

Automated radio telemetry systems (ARTS) have the potential to revolutionise our understanding of animal movement by providing a near-continuous record of individual locations in the wild. However, localisation error in data generated by ARTS can be very high, especially in natural landscapes with complex vegetation structure and topography. This curtails the ecological questions that may be addressed with this technology. Here, we set up an ARTS grid in a valley with heterogeneous vegetation cover in the Colombian high Andes and applied an analytical pipeline to test the effectiveness of localisation methods. We performed calibration trials to simulate animal movement in high-or low-flight, or walking on the ground, and compared workflows with varying decisions related to signal cleaning, selection, smoothing, and interpretation, along with four multilateration approaches. We also quantified the influence of spatial features on the system’s accuracy. We tested the grid by deploying tags on two high-altitude hummingbirds, the Great Sapphirewing (*Pterophanes cyanopterus*) and Bronze-tailed Thornbill (*Chalcostigma heteropogon*). Results showed large variation in localisation error, ranging from only 0.4–43.4 m from known locations up to 474–1929 m, depending on the localisation method used. The lowest average median error across calibration tracks was 105 m. In particular, we found that the selection of higher radio signal strengths and data smoothing based on the temporal autocorrelation in movement data are useful tools to improve accuracy. Moreover, the variables that significantly influence localisation error include terrain ruggedness, height of movement, vegetation type, and the location of animals inside or outside the grid area. In the case of our study system, thousands of location points were successfully estimated for two hummingbird species that previously lacked movement ecology data. Our case study on hummingbirds suggests ARTS grids can be used to estimate small animals’ home ranges, associations with vegetation types, and seasonality in occurrence. We present a comparative localisation pipeline, highlighting the variety of possible decisions while processing radio signal data. Overall, this study provides guidance to improve the resolution of location estimates, broadening the application of this tracking technology in the study of the spatial ecology of wild populations.

## Introduction

Tracking animals has been a revolutionary tool in ecology and evolution, since the movement of animals determines the spatiotemporal patterns of species assemblages, and thus affects ecological processes and ecosystem functioning (Kays et al., 2015). Movement data can improve our understanding of fundamental questions regarding species’ habitat associations, home range sizes, behaviour, dispersal, and migration (Hansson & Åkesson, 2014). In the face of ongoing global change, a better understanding of animal movement is crucial to guide effective conservation strategies (Runge et al., 2014) and protect threatened wildlife (Wilcove & Wikelski, 2008). Yet even with the rapid advancement and miniaturisation of animal tracking technologies in recent years, which has significantly increased the range of animals that can be tracked (Wilmers et al., 2015), we still lack essential movement data for many small species under 20 g.

The technologies most frequently used to track terrestrial animals with small body size include global positioning systems (GPS), geolocators, passive integrated transponders (PIT) paired with radio frequency identification (RFID) antennas, and radio telemetry tracking with handheld antennas or automated receivers (Bridge et al., 2011). GPS tags communicate with satellites to determine longitude, latitude, and time; however, the lightest tags using GPS technology are archival, which do not transmit data remotely but must be recovered to retrieve data stored on-board. Another limitation of GPS tags is their size, with very light archival GPS tags still weighing over 0.95 g (e.g., Pathtrack, UK: https://www.pathtrack.co.uk/). Geolocators are lighter (∼0.35 g) but also need to be recovered to download archived data. They have been mainly used for long-distance movement, since they depend on changes in sunlight across latitudes to infer location and thus are more suitable for studies over large spatial scales (McKinnon & Love, 2018).

To determine a location of an animal at finer spatial resolutions, PIT tags provide a useful alternative: these tags emit radio waves with an unique individual ID when near a receiver (Gibbons & Andrews, 2004) and are light enough to be used on animals with a body mass of around 2 g (e.g., D’Arcy et al., 2020). The main drawback of PIT tags is that animals have to be very close to RFID antennas for detection to occur (though the detection distance depends on frequency, tag size, and tag orientation, it is generally under ∼30 cm [Ousterhout & Semlitsch, 2014]). As such, PIT tags are mostly used to record the presence or absence of an individual rather than to infer an animal’s trajectory. In addition, to ensure detection, the associated receivers are usually baited (such as antennas equipped on hummingbird feeders, see for example Falk et al., 2021), possibly influencing animal behaviour and thus problematic for some research questions.

Very high and ultra high-frequency (VHF and UHF) radio telemetry uses radio waves that are detected by receivers. The lightest radio transmitters are between 0.6 and 0.15 g (e.g., Cellular Tracking Technologies - CTT, USA, celltracktech.com; Lotek Wireless INC, UK, lotek.com) and may be used on very small animals such as insects (Kissling et al., 2013). Animals with radio tags have been most frequently tracked by handheld antennas, generally limiting those studies to comparatively small areas and short or non-continuous time periods. In addition, very rugged terrain is difficult to traverse or sometimes cannot be accessed at all (e.g. Hazlehurst & Karubian, 2018) and animals may possibly change behaviour in the presence of researchers carrying an antenna. However, automated radio telemetry systems (ARTS) have overcome these limitations by continuously scanning for radio signals from transmitters and identifying more than one transmitter at the same time (Kays et al., 2011). Single antennas are used to identify the presence of an individual in the area (e.g. Motus; Taylor et al., 2017), while arrays of receivers can be used to estimate location based on the relationship between recorded radio signal strength (RSS, measured in decibels) and distance from each receiver (Luomala & Hakala, 2022). The distances around receivers are radii whose overlapping circumferences may be calculated through triangulation (e.g. Ward et al., 2013) or multilateration to give a location estimate (e.g. Paxton et al., 2022).

However, in natural landscapes, the overlap of circumferences around nodes will not be perfect, because vegetation and topography modify or obstruct radio signals, adding noise to the relationship between RSS and distance and thus increasing error of localisation estimates (Ward et al., 2013). In addition, radio signal strength may vary with the orientation of the transmitter’s antenna in relation to the receiver (Fisher et al., 2021), the strength of the transmitter (Ward et al., 2013), and the receiving capacity of the receiver, as well as with the movement of the transmitter itself (Lenske & Nocera, 2018) when an animal is following a trajectory path. The optimization of an overlapping location may be achieved through non-linear least squares (NLS) regression (as in Paxton et al., 2022). Analytical efforts to reduce error have included using Bayesian approaches to calculate parameters of the RSS-distance relationship (Wallace et al., 2022), constructing kernel density estimates around receiving nodes (Wallace et al., 2022), filtering cut-off thresholds based on signal strength or distance (Celis-Murillo et al., 2016, Paxton et al., 2022), and incorporating k-means clustering to remove outliers (Luo et al., 2022 apply it in an indoor setting).

The variety and relative novelty of approaches makes choosing a single one difficult when setting up new ARTS grids and warrants more testing of grids in different configurations and landscapes. Moreover, recent studies that report substantial reductions in error have been limited to simple landscapes with little to no vegetation and flat topography (e.g. Paxton et al., 2022) or grids that cover very small areas (e.g. Wallace et al., 2022). Although such studies provide useful guidelines, broadening the assessment of localisation methods to more complex landscapes is necessary. Without a better understanding of the localisation error and limitations of ARTS grids, it is difficult for researchers to decide what tracking technology to use in light of their questions of interest.

We tested an ARTS grid in a natural landscape in the high Andes with moving calibration trials simulating flight at two heights and movement on the ground in order to assess the suitability of the grid to collect movement data on small animals. Although ARTS grids are mostly calibrated with stationary tests (Fleming et al. 2020, but see Mennill et al., 2012, Krull et al., 2018, Fisher et al., 2021), our approach more closely resembles how radio signals are affected by continuous motion of tracked animals. In addition, the inherent autocorrelation of movement tracks may provide tools to reduce error through time series smoothing such as splines or state-space models (Newman et al., 2023). We set up a data processing workflow for localisation and compared different decision pathways in the steps of signal selection, smoothing, interpretation and multilateration. We also tested how localisation error is influenced by different spatial features of the landscape and the locomotion type of study species.

This work evaluates the feasibility and accuracy of automated tracking of small animals in natural landscapes, using high-Andean ecosystems as the location of our ARTS grid. Small animals we have targeted to tag in this area include species of birds, amphibians, reptiles, rodents, and bats. Across taxa, species in these habitats are highly endemic (Flantua et al., 2019, Sonne et al., 2022), increasingly threatened by changes in climate and land use (Cresso et al., 2020, Valencia et al., 2020), and strikingly understudied in terms of their movement patterns (Jahn et al., 2020). Furthermore, small animals present strong limitations on the weight they can safely carry (e.g. precluding the use of GPS trackers) and most attempts to track fast-moving animals, such as hummingbirds, have been limited to hand-tracking, limiting the studies to a few focal individuals (e.g. Pavan et al., 2020). We tested our ARTS grid by tagging two hummingbird species and collecting basic information on their movement trajectories and utilisation distributions.

## Methods

### Automated radio telemetry grid

The installed ARTS grid is composed of 46 receiving nodes (CTT Node V2, Cellular Tracking Technologies, USA) and a base station (CTT SensorStation) set in the Valle de los Frailejones inside Chingaza National Natural Park, Colombia (4°31’30.5‘N 73°46’23.9’W). Each node continuously receives unique tag IDs and radio signal strengths (RSS, measured in decibels) from tags emitting radio signals at 433 MHz. Nodes are set in the valley on steel conduits (electrical metallic tubing) approximately at 2.5 m above the ground and 150 m apart. Nodes send collected tag information, RSS, and a timestamp to the base station, which has four receiving Yagi antennas (∼10 m above the ground) and is connected to a deep-cycle marine battery and a solar panel for power (Figure S1). Data is downloaded manually from the base station since there is no available internet connection in the area. The grid of receiving nodes covers an area of 0.72 km^2^. It is situated in a valley where ground elevation varies between 3162 and 3391 m asl (Figure S2) and is covered by natural vegetation consisting of paramo rosetted plants (*Espeletia grandiflora* and *E. argentea*) and shrubs (*Pentacalia* spp.), elfin forest, bamboo (*Chusquea tessellata*), and grassland (Figure 1 and Figure S3).

**Figure 1.**
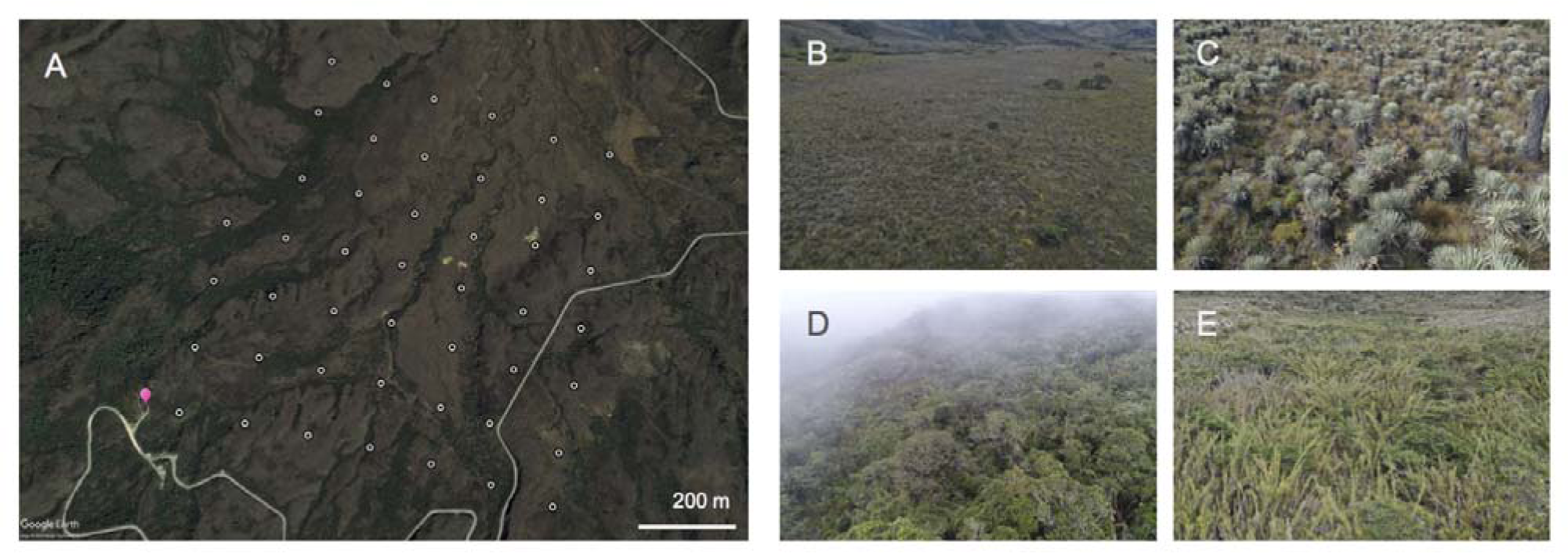
A) Automated radio telemetry grid installed in the Valle de los Frailejones, Chingaza National Natural Park, Colombia. Each of the 46 receiving nodes installed in the valley are represented by a white circle, and the central antenna is indicated with a pink marker. Satellite image reproduced in Google Earth Pro. The main vegetation covers inside the grid include B) open grassland and flooded areas, C) paramo rosetted plants and shrubs, D) elfin forest, and E) bamboo. Photographs taken with DJI Mavic 2 drone by Carolina Arévalo (B and C) and Nicolás P. Skillings (D and E).

### Calibration trials

To simulate animal movement at different heights above the ground, radio transmitters were tested within the grid with moving trials, either attached to a flying drone (DJI Mavic 2) or at the end of a wooden stick while walking. Researchers flying the drones or walking were asked to randomly move within the grid, and the trials were distributed in such a way that most of the area of the grid was covered (see Figure 5D). Drone flights were on average 39 (± 16 sd) m above the ground, and walking trials at waist level (∼1 m agl) or ground level (∼ 0 m agl), thereby imitating movement of flights above the height of vegetation, at vegetation height, and walking at ground level, respectively. We used 19 tags, of which 3 were PowerTags (CTT PowerTags, Cellular Tracking Technologies, USA) and 16 LifeTags (CTT LifeTags, Cellular Tracking Technologies, USA). PowerTags depend on an internal battery for energy, and radio signals were set to emit every 60 seconds. LifeTags use miniaturised solar panels and emit a signal every 2 seconds. Of the 16 LifeTags we used, 11 were designed for hummingbirds with a 3D-printed chassis (design included as supplementary material) and five were intended for larger birds with the attachment method that comes from the manufacturer.

We carried out calibration trials between September 26th and November 7th 2022 and 6:00 and 20:30 local time, with some nocturnal trials to test PowerTags that do not depend on solar energy. Before starting a trial, we confirmed that each tag was functional using a handheld antenna (CTT Locator). The locations and timestamp of tags during the trials (relocations) were recorded with the incorporated drone GPS for flights and with a handheld GPS (Garmin eTrex 32X) during walking trials. Drone tracks were cleaned to remove initial straight-line flights that had the purpose of reaching more distant parts of the grid.

### Estimating location from radio signal strength

RSS values that were recorded by receiving nodes were downloaded from the main antenna, and processed in a data workflow wherein different pipelines were explored to compare how decisions in the steps of signal selection, interpretation, smoothing, and localisation would affect error (Figure 2). First, we filtered signal data to discard noise by retrieving only the signals belonging to the tested tag during calibration trials. Then, we matched signals to the nearest timestamp of known relocations during calibration, within a maximum difference of 10 s to allow for delays in signal reception and possible mismatches between GPS and node clocks (Figure S4). For each relocation and node, the maximum and average RSS was calculated to account for radio signal bouncing and multipathing (Paxton et al., 2022). Average RSS values were calculated after removing signals that were 4.33 dB or more away from the calculated average value and therefore considered to be outliers (since 95% of signals were within 4.33 dB of average RSS).

**Figure 2.**
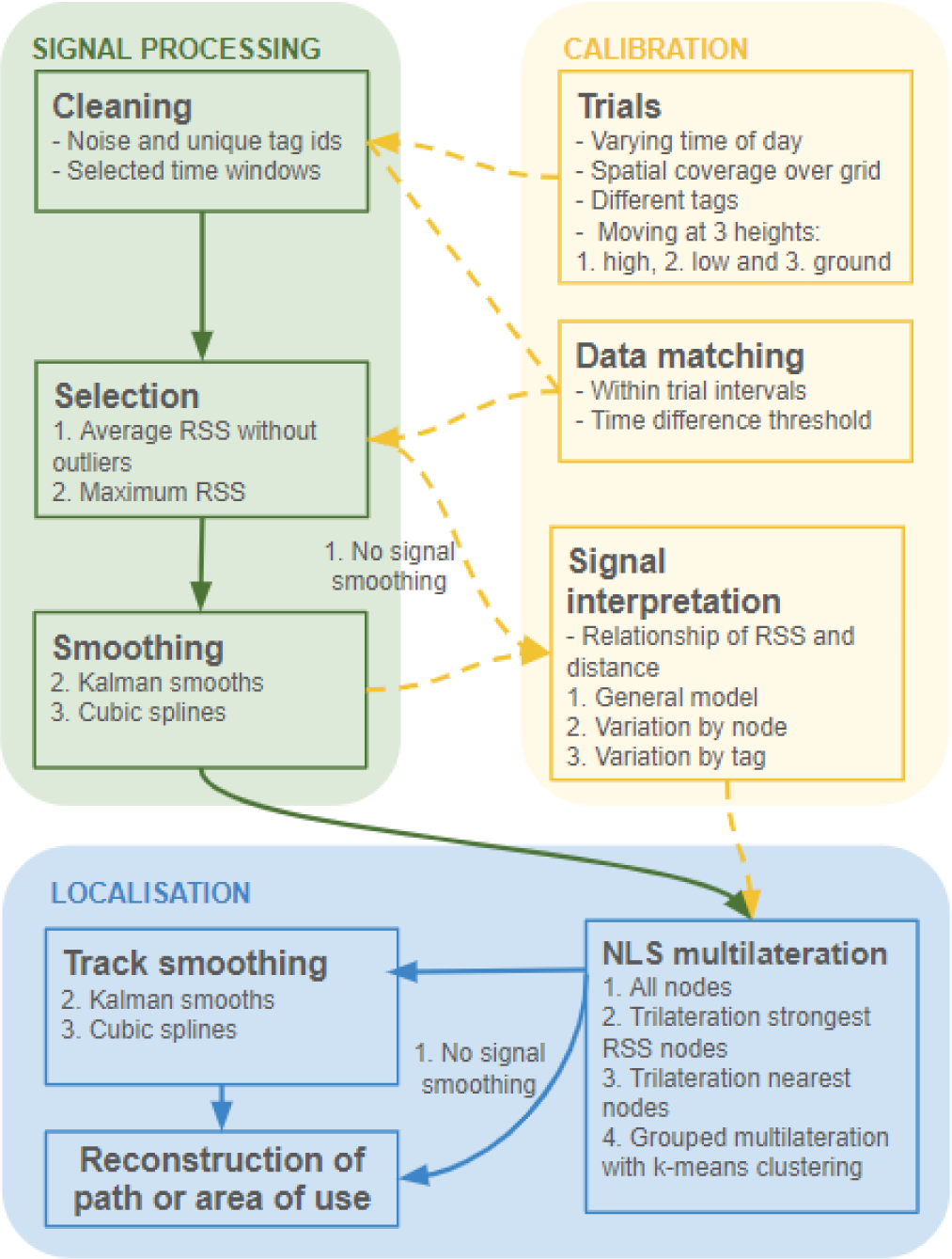
Workflow to estimate locations from received radio signal strengths, by processing and calibrating signals for localisation. Once the system is calibrated and animals are being tracked, the sequence of steps may be followed along the solid lines, excluding the dashed lines that indicate calibration. In each step, the different approaches explored here are shown in a numbered list. Exploring the different decision pathways results in 216 repeated measures of error for each relocation within high flying (drone), low flying (walking with tag at 1.5 m), or ground (walking with tag at 0 m) trials.

After the selection of average and maximum RSS reads, we added a signal smoothing step using either cubic splines or Kalman smoothers. This step corrected outlying values and reduced noise in the relationship between RSS and distance by using the implicit temporal autocorrelation of tracks, wherein the RSS value at one time step is influenced by the value in the previous and future time step. Cubic splines were calculated with the function “smooth.spline” available in R (R Core Team, 2023), using generalised cross-validation to find the optimal model parameters. To apply a Kalman smoother, we used the package “KFAS” in R (Helske, 2017). We fitted a state-space model by individual calibration track and node, using equal time intervals and a local-level model. Model parameters were initially set as unknown and then estimated by maximum likelihood using an unconstrained non-linear optimization method “BFGS” incorporated in the function “fitSSM” of the “KFAS” package. Time series of nodes that had less than 5 or 2 consecutive reads could not be smoothed with cubic splines or Kalman smoothers, respectively, so their initial and non-smoothed values were used instead.

The exponential decay relationship between maximum or average RSS and distance was modelled in a two-step approach in which we first fit an NLS function to the general dataset and then updated the model estimates to allow them to vary independently by node or tag. To do so, we calculated the distance between known relocations and each node in metres to obtain true distances and used the following equation as in Paxton et al. (2022):

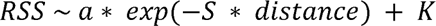

where *a* is the intercept, *S* is the decay factor, and *K* is the horizontal asymptote in the exponential relationship. To find initial parameters *a*, *S,* and *K*, we employed the self-starting function “SSasymp” in the “nls” function in R and used resulting estimates as the values for the final general model (Paxton et al., 2022). Then, the final general model was used to fit models by node or tag. Allowing parameters to vary by node is important to convey variation of not only the detection capacities inherent in the receivers themselves (Harbicht et al., 2017) but also the different environmental characteristics that surround each node. Variation of parameters between tags may be due to differences between the transmitters caused by imperfections during manufacture. Nodes with models that did not converge were allowed to return to the final general NLS model estimates.

Estimated distances from fitted models were then used in four different localisation methods: 1) multilateration using signal reads from all the nodes that detected transmitters at each relocation, 2) trilateration with only the three nodes with strongest signal, 3) trilateration with three nearest nodes in estimated distance to relocation, and 4) grouped multilateration with k-means clustering. The fourth method consists of producing multiple location estimates per relocation by using all the possible combinations of subgroups of nodes that detected a signal (where group sizes must be of 3 nodes or more). Due to computing limitations, we set the maximum number of nodes to use to 7 and selected those with strongest signal reads (Supplementary methods). We clustered resulting estimates in k-groups to reduce the sum of squares within clusters (Hartigan & Wong, 1979). The optimal value of k may be found as the total within-cluster sum of squares is reduced. In this way, extreme or spatially biassed values will be filtered and the centroid of the largest cluster may be selected as the final estimated location (Luo et al., 2022).

For all methods, relocations that were detected by less than 3 nodes were excluded, as well as signal reads below the estimated horizontal asymptote. Negative estimated distances were changed to a distance of 0 m, being assumed to be very close to the node.

### Factors that influence error

Error, as a measure of accuracy, was calculated as the difference in metres between known and estimated locations. Given that we estimated locations through different decision pathways (Figure 2), each relocation of calibration trials had repeated measures of error (216 possible combinations). To compare error between measures and identify how analytical decisions influence accuracy, we fit a linear mixed effects model (LMM) with each relocation nested within its calibration trial as the random effect and all steps in data processing as fixed effects. Models with all the possible combinations of fixed effects were compared and ranked by Akaike Information Criterion corrected for small sample sizes (AICc, Supplementary methods). In addition, we generated random tracks for each calibration trial (99 simulations for each) to compare localisation errors between estimated tracks and what would be expected from a random process, using continuous-time movement models fitted with the package “ctmm” (Calabrese et al., 2016, Fleming & Calabrese, 2023) and handling trajectories with packages “move” (Kranstauber et al., 2023) and “trajr” (MacLean & Showron Volponi, 2018) in R.

After selecting a single method to estimate locations, we also tested for the influence of spatial features on error. For this, we used a LMM with calculated error as response variable, calibration trial as random effect, and ground elevation, terrain ruggedness, distance to the edge of the grid, presence inside or outside the grid, flight height, and dominant vegetation cover type, as well as all the interactions between vegetation cover and the first five landscape variables, as fixed effects. However, ground elevation and terrain ruggedness were correlated (Pearson correlation, r = 0.819, p < 0.05), so we chose to only include the latter in the models (Supplementary methods).

Data on ground elevation was downloaded from the GLO-30 Copernicus digital elevation model (30 m resolution) of the European Space Agency (ESA, 2024). Terrain ruggedness was calculated as the standard deviation of ground elevation values in a 100 m radius buffer around each known relocation. Flight heights were estimated as the difference between elevation reads from recording GPS devices (handheld or drone) and ground elevation. Vegetation covers were mapped with hand-drawn polygons on Google Earth Pro using satellite imagery and extensive on-the-ground knowledge of the valley to detect differences between four main categories of open grassland, typical paramo vegetation of rosetted plants and shrubs, dense vegetation of either elfin forest or bamboo, and built-up areas pertaining to a dirt road and park rangers’ housing. Forest and bamboo were lumped into a single category because they were not distinguishable from Google Earth imagery and likely attenuate radio signals similarly due to their heights and dense structure. Polygons were then rasterized into a 1 m resolution grid in R using packages “sf” (Pebesma & Bivand, 2023) and “raster” (Hijmans, 2023). The proportion of each vegetation type was measured from the covered area within a 100 m radius, as the dominant vegetation type around a relocation was defined as the one with the highest occupied area.

Autocorrelation structure in residuals was assessed for all LMMs (Figures S6, S7, S9). We used the packages “nlme” (Pinheiro & Bates, 2023) and “MuMIn” (Bartoń, 2023) in R to fit and evaluate LMMs.

### Animal tracking

Hummingbirds were captured using mist nets (32 mm mesh, 2.5 x 6, 9, or 12 m) that were open during the morning (6–11 AM) and checked in 15–20 minute net rounds depending on the weather. Captures occurred between December 2022 and February 2023, for a total sampling effort of 207 net-hours in total (1 net-hour = 1 h of one 12 m net, Table S1). Handling never lasted more than 10 minutes, and all birds were fed sugar water prior to release (Russell et al., 2019). Animals were handled by trained experts with research and ethics permits (Pontificia Universidad Javeriana #127-22, University of Washington IACUC protocol number: 4498-05, and University of Aberdeen approval 01/19/22). Research in Chingaza National Natural Park was carried out under the permit #20212000005863.

We deployed tags on 10 adult individuals of the two larger hummingbird species of the area: 3 male and 1 female Great Sapphirewing (*Pterophanes cyanopterus*) and 6 male Bronze-tailed Thornbills (*Chalcostigma heteropogon*). LifeTags and harnesses weighed 0.45 g, while PowerTags weighed 0.35 g, within ∼3-5% of body mass (Table S2). LifeTags were attached on Great Sapphirewings using 0.5 or 0.7 mm StretchMagic string harnesses as designed by Williamson and Witt (2021), which have been successfully used to track migratory Giant Hummingbirds (*Patagona gigas*), remaining on birds for several years without any visible effect after long-distance migration. Powertags and LifeTags were attached on back Bronze-tailed Thornbills using non-irritating superglue (Loctite PureGel) to attach tags on scapular area (Hadley & Betts, 2009).

Before releasing tagged hummingbirds, 10-minute flight trials were conducted inside a closed tent (330 x 220 x 115 cm) with a central perch made from string. The trials were intended to evaluate the influence of tags on mobility before release, and therefore whether birds could fly, hover, and perch. All tagged hummingbirds were able to do all of these activities with no observable change in behaviour, so they were released, and at least two people observed subsequent flight and perching behaviour with binoculars. Continued monitoring with mist-netting allowed us to also evaluate effects of tags on recaptured birds by recording body condition and noting any signs of wear or abrasion.

Hummingbird tracks were calculated using the steps of the processing workflow that led to lower error in LMMs assessing the influence of analytical decisions. Resulting track coordinates were cleaned by filtering out positions with outlying speeds that would be unfeasible even in high-speed-flapping flight (Chai et al., 1999 report 14.4 m/s for Ruby-throated Hummingbirds) or courtship dives (Clark, 2009 reports 27.3 m/s for Anna’s Hummingbirds). We do not know of studies that have calculated power flight curves to estimate maximum horizontal flight speeds (Engel et al., 2010) for our tagged species. Given that our knowledge about hummingbird flight speeds in general is scarce, we chose a maximum speed threshold based on the distribution of speeds resulting from estimated locations. Utilisation distributions were estimated with autocorrelated kernel densities (AKDE) through continuous-time movement models (Silva et al., 2022) that incorporate measures of error and allow for unequal time intervals in the R package “ctmm” (Fleming & Calabrese, 2023).

## Results

### Propagation of signal

Calibration trial tracks had an average time difference between relocation points of 2 (± 3) s for drone trials and 3 (± 2) s for walking trials (Figure S4). Individual trials lasted on average 389 (± 99), 261 (± 54) and 371(± 139) s, covered distances of 1069 (± 416), 111 (± 45) and 106 (± 64) m and moved at speeds of 3.57 (± 1.78), 0.44 (± 0.12) and 0.31 (± 0.06) m/s, for drone flights, mid level and ground level walking trials, respectively.

From the 77 calibration trials we carried out, 60 trials (15 high flight, 22 low flight, and 23 ground-level trials) were detected by the ARTS grid, with 43,323 radio signal reads for 2,361 relocations in total. Trials that were performed close to sunset (after 16:30) were not detected by receivers at all (10 ground and 3 high flight trials), possibly because light levels were not sufficient for LifeTags to emit a signal. Out of the 19 tested tags, 3 (2 LifeTags and 1 PowerTag) were not detected at all by the grid during calibration, even though they were detected by a handheld antenna to test that the tags were working before trials began. The median value for time intervals between signal emissions was 61 s for PowerTags and 5 s for LifeTags. The number of receiving nodes recording a signal differed between the three test heights (log likelihood = −7895.1 [7889.5 null], chisq = 11.246, p = 0.003), with fewer nodes recording a signal for ground-level trials on average (12 [± 8] nodes) compared to flying trials (14 [± 9] and 14 [± 7] nodes for drone flights and mid-level walking trials, respectively), although estimated confidence intervals were not different from zero (Table S3). One node failed during calibration trials and did not record any signals. Recorded signal strength varied between −37 and −115 dB, with 95% of recorded signals between −79 and −110 dB (Figure 3).

**Figure 3.**
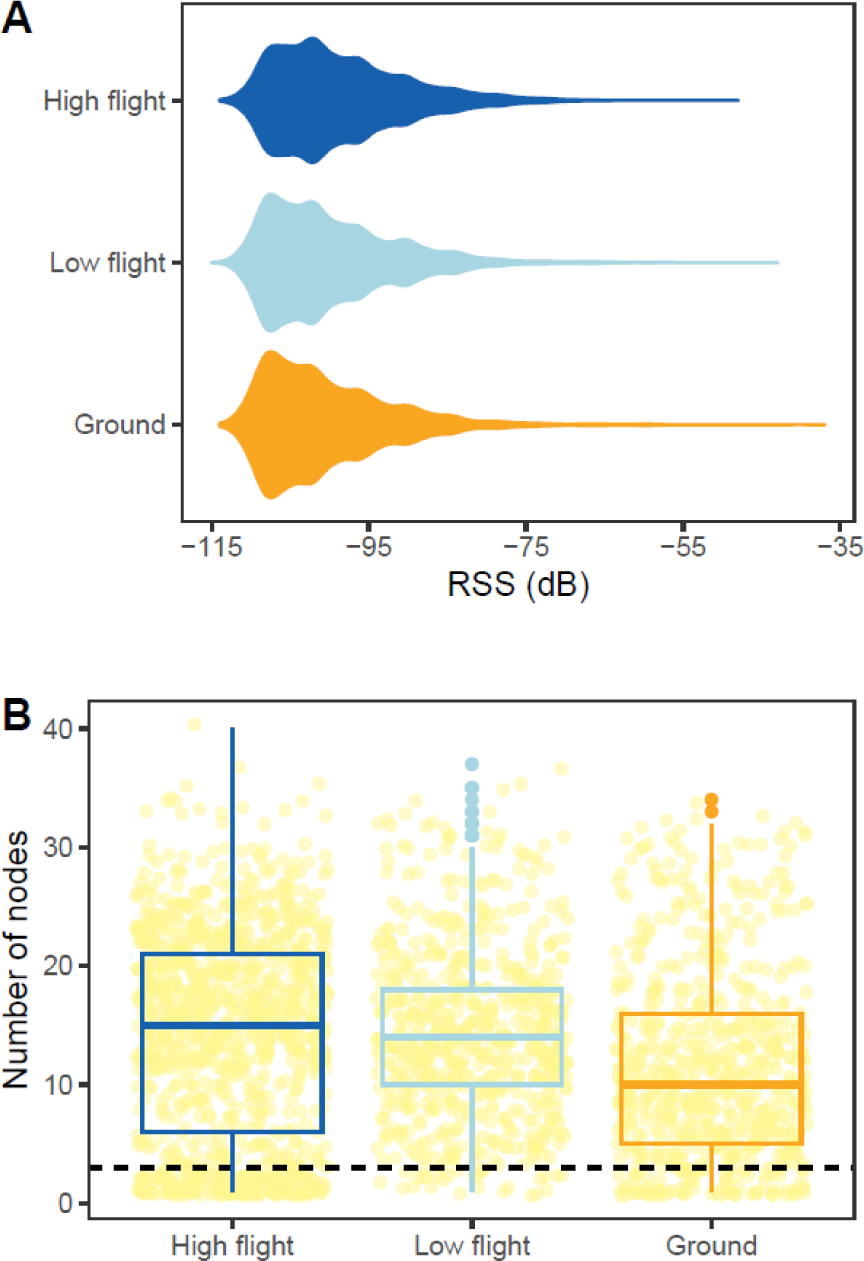
A) Distribution of detected radio signal strength (RSS) measured in decibels (dB) when calibration trials were carried out at high flight (dark blue), low flight (light blue) or near the ground (orange). B) Number of nodes that detected a relocation, shown as a yellow point. Boxplots show median values by height of calibration trials with middle thick line, first and third quartiles with box hinges and 1.5*interquartile range with whiskers. Horizontal dashed line indicates 3 nodes, under which localisation is not possible.

We found that signal strength decreases with distance (Figure 4) and may be modelled with an exponential decay curve. Standard error of residuals decreased when RSS was smoothed with cubic splines and Kalman smoothers and when models were fitted independently by receiving nodes or tags (Figure 4). Although residuals in exponential decay models showed strong autocorrelation (Figure S6), incorporating an autocorrelation structure to models led to singular results. However, robustness of estimated model parameters was shown by small 95% confidence intervals and parameter standard errors calculated through bootstrapping with 999 iterations (Table S4), so models were used to calculate estimated distances of signal to nodes.

**Figure 4.**
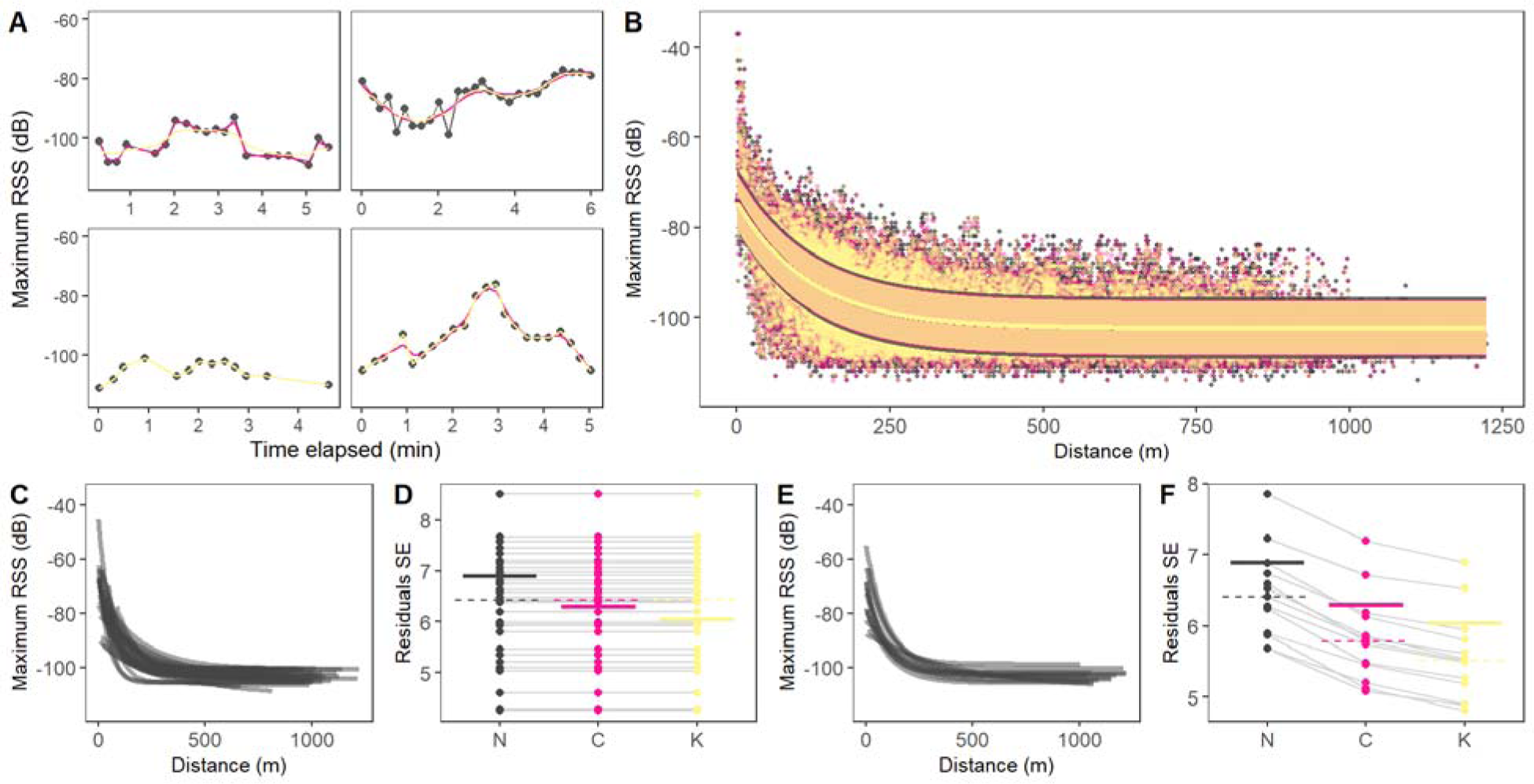
Detected maximum radio signal strength (RSS) and variation of fitted exponential decay models. Colours throughout all panels represent maximum RSS values that were not smoothed (black), smoothed with cubic splines (pink), or Kalman smoothers (yellow). A) Example of maximum RSS detected for one calibration trial by four different receiving nodes through time. Points show initial reads and lines indicate smoothed values with cubic splines or Kalman smoothers. B) Relationship between maximum RSS and measured distance for calibration trials. Line shows fitted non-linear least squares exponential decay models and shaded area indicates standard error (SE) of residuals. Models fitted individually by receiving C) nodes or E) tags; each line represents a separate node or tag. Standard error of residuals in models by D) node or F) tag, for maximum RSS values that were not smoothed (N), smoothed with cubic splines (C), or Kalman smoothers (K). Points show individual receiving nodes or tags, and lines connect the same node or tag. Solid thick horizontal lines show SE for the general model and dashed horizontal lines show median value for models separated by nodes or tags. See Figure S5 for graphs of average RSS.

### Localisation error

All methods generated localisation errors that were significantly better than a random process (paired Wilcoxon rank test, p < 0.05 Table S9). Depending on the localisation approach, some relocations had estimates that were only a few metres away (recorded minimum 0.4 m and up to 43.4 m depending on the method, Table S10) from the true location but the variation in localisation error within trials was high (average standard deviation 99 [± 61] m across methods) and maximum error values reached 473.95–1928.91 m. Error was greatest when trilateration was performed by selecting the nearest nodes (Figure 4, Table S10), and although all steps in the localisation workflow influence accuracy (top-ranking model included all predictors with 99.99 % AICc weight, Table S11), the method of selecting the nearest nodes during trilateration had the largest effect magnitude in the models (Figure 5). Selecting maximum RSS, smoothing signals and tracks, using general model parameters, and trilateration with the three strongest signal nodes are choices that will lead to reduction in localisation error (Table S12). The method with the lowest error resulted in an average median value of 105 m (Table S10, high flight, maximum RSS, cubic splines smoothing and node-specific model parameters). However, location estimates of more relocations were possible when decay model parameters were allowed to vary by nodes, since most had a lower horizontal asymptote than the general models (Table S6)

**Figure 5.**
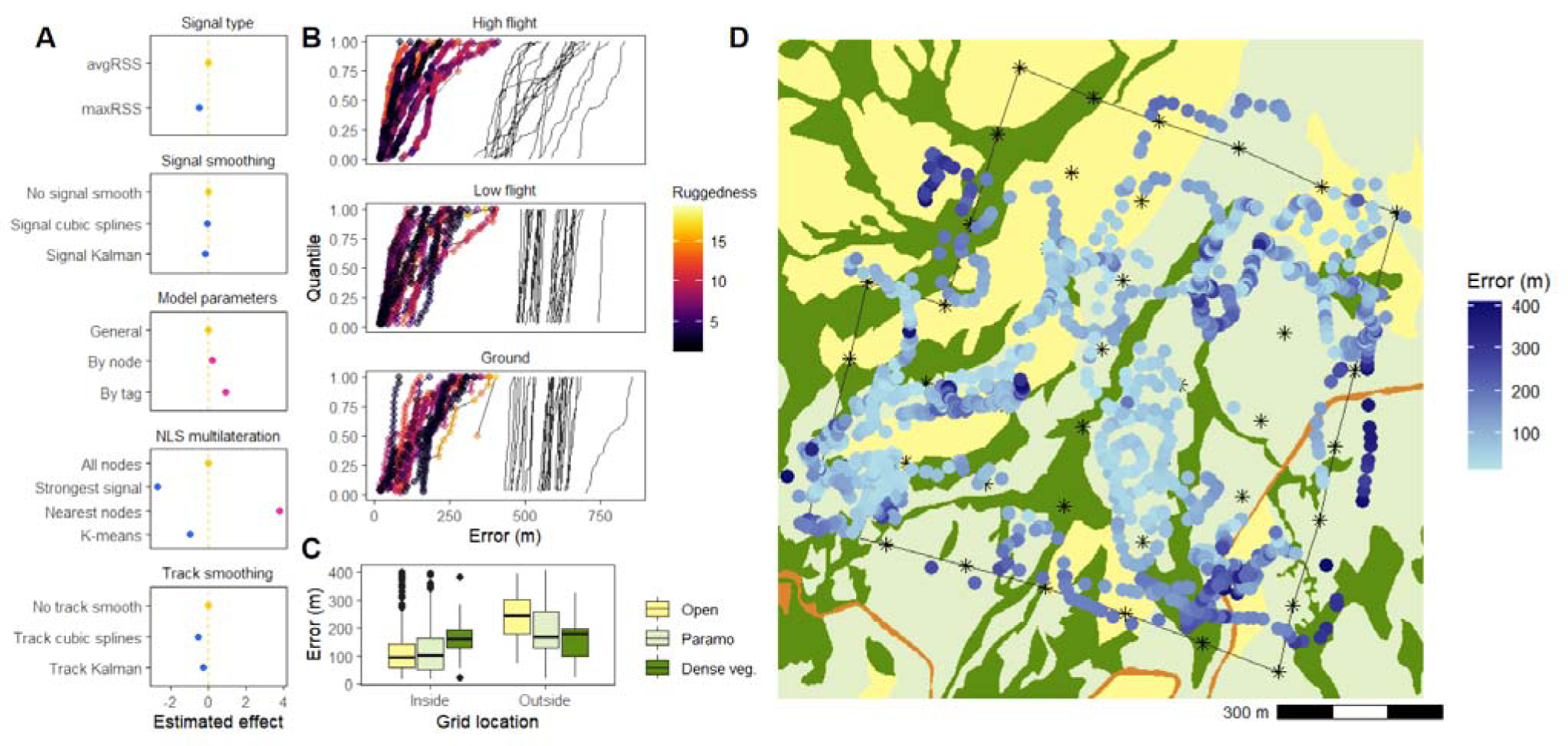
A) Estimated effects of different decision steps on methodological workflow on localisation error, measured as the distance in metres between known and estimated locations. Panels represent steps of signal type selection, smoothing, signal-distance decay model parameters, non-linear least squares (NLS) multilateration, and track smoothing. First row in each panel is the baseline method (Table S12) for model comparison (thus has an effect of 0, indicated by yellow colour and vertical dashed line) and blue and pink show reduced or increased error, respectively. Points are estimated effects and horizontal lines 95% confidence intervals (barely visible because calculated intervals were small). Effect magnitude shown has the square root transformation to keep model assumptions of normality. B) Quantiles of calculated error in tracks of high flying (drone), low flying (walking with tag at 1.5 m) or ground (walking with tag at 0 m) calibration trials. Points show relocations and are connected with lines to show tracks. Colour gradient represents terrain ruggedness in metres around 100 m radius of relocation. Solid black lines show error calculated for simulated tracks (99 iterations for each) of a random localisation process. C) Error for relocations inside and outside of the automated radio telemetry (ARTS) grid, according to dominant vegetation type in a 100 m radius. Vegetation types are classed as open grassland, paramo, and dense vegetation (forest and bamboo). D) Map of error calculated from calibration tracks (blue colour scale), with receiving nodes shown as asterisks and the edge of the ARTS grid as a black line. Colours of vegetation types are the same as panel C, and orange colour represents a dirt road.

For the effects of spatial features on error, we found that dominant vegetation type, flight height, terrain ruggedness and whether a relocation was inside or outside of the grid all had an effect on error that was different from zero (Figure S10, Table S14). The largest effect magnitude was related to location in or out of the grid, with relocations outside substantially increasing in error (36–9-fold with 95% CI). Contrary to our expectations, error decreased in both dense and paramo vegetation compared to open grassland. However, this might be due to increased error of relocations outside of the grid (Figure 5, Figure S10). Inside of the grid, relocations that were located either in paramo or dense vegetation increased error by 2 (0.01–11 with 95% CI) or 22 (5–52 with 95% CI) times, respectively (Table S14), compared to open grassland. No relocation had a dominant vegetation cover of built-up area, so this type of vegetation cover was not included in the models.

### Tracked hummingbirds

Signals were detected for 120,989 relocations of tagged hummingbirds (Figure 6, Figure S11 and S12), with 56,840 (47%) and 43,774 (36%) being detected by at least 3 nodes with RSS reads over the horizontal asymptote when using node-specific and general model parameters, respectively. One Bronze-tailed Thornbill had no relocations that could be estimated because all recorded RSS values were −103 dB or less and therefore under the horizontal asymptote of the RSS-distance exponential decay model. Another recaptured Bronze-tailed Thornbill lost its tag on the day following attachment, even though it was not moulting any feathers, suggesting that the glue-on method is not appropriate for longer-term tracking. In contrast, tags attached with the harness method showed potential for longer-term tracking. One female Great Sapphirewing was recaptured 34 days after tag attachment and showed good body condition (body mass, pectoral muscle, and feather wear) and no signs of abrasion, with the attached tag still having a good fit and no deterioration. As evidenced through direct observation with binoculars, all hummingbirds flew normally and perched on flowers to continue foraging after release. Although two Great Sapphirewings appeared to have left the area during the period they were tracked, another two remained in the grid and generated tracking data for multiple weeks (for 43 and 71 days, Figure 6E). Minimum time intervals between detections were 61 s for Powertags and 5 s for Lifetags.

**Figure 6.**
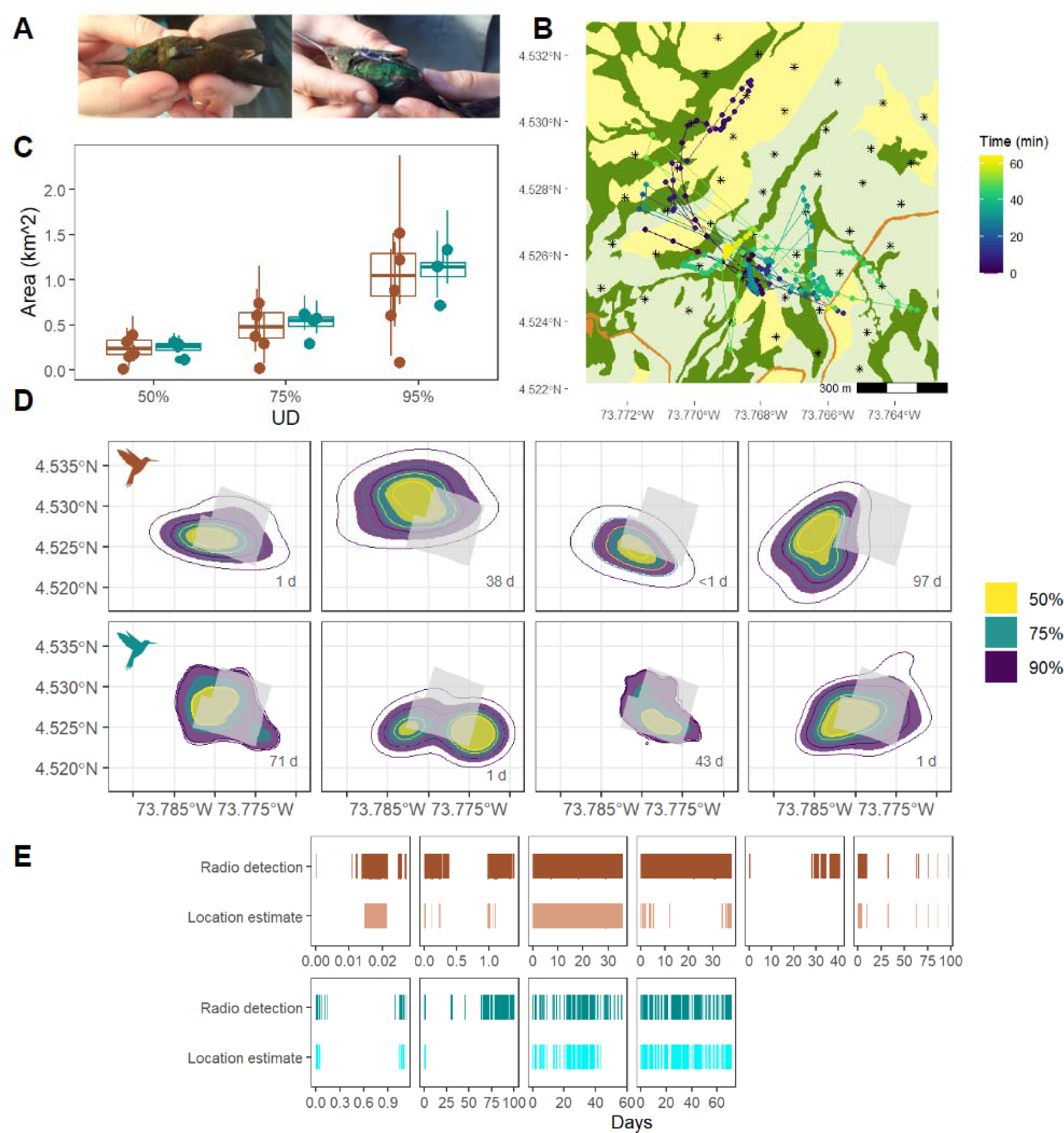
Tracking data from the Chingaza automated radio telemetry grid reveals movement trajectories and utilisation distributions for two hummingbird species in a high-Andean landscape. A) Tagged Bronze-tailed Thornbill (*Chalcostigma heteropogon*) with glued on PowerTag (Cellular Tracking Technologies - CTT) and Great Sapphirewing (*Pterophanes cyanopterus*) with harness LifeTag (CTT), from left to right. B) Example track of a male Great Sapphirewing during the first hour of the morning (242 fixes, from 6:55 to 8:00 local time); relocations are represented with points and consecutive points in time connected with lines to show path trajectory. Colour of points indicates passing time in minutes; vegetation covers of open grassland, paramo and dense vegetation are coloured in yellow, light green and dark green, respectively. Orange colour shows the dirt road. C) Estimated area of utilisation distributions (UD) with 50, 75 and 95% autocorrelated kernel density estimates (AKDE) for tracked Bronze-tailed Thornbills (brown colour) and Great Sapphirewings (turquoise). Points show individuals and vertical line ranges indicate 95% confidence intervals. Boxplots show median values by species with middle thick line, first and third quartiles with box hinges and 1.5*interquartile range with whiskers. They do not include one Bronze-tailed Thornbill with a very small UD that was considered an outlier (shown as lowest brown point). D) AKDE for 8 hummingbirds (top row Bronze-tailed Thornbills, bottom row Great Sapphirewings). The outlier Bronze-tailed Thornbill with very small UD is shown in Figure S13. Filled polygons show estimates for 50, 75, and 95% AKDEs, with lines indicating lower (inner line) and upper (outer line) 95% CI for each. Numbers in bottom right corners of panels are the time periods over which each individual was tracked. Light grey polygon indicates the area covered by the automated radio telemetry grid. E) Signal detection and location estimates through time, shown as days, for tagged hummingbirds (top row Bronze-tailed Thornbills, bottom row Great Sapphirewings).

To estimate locations of tracked hummingbirds, we used maximum RSS, cubic splines smoothing of signals and tracks, node model parameters, and strongest signal localisation, which had the lowest average median error of 105 m. Although models indicated that changing RSS-distance model parameters to vary by node or tag increased error, the magnitude of this effect was low (Figure 5, Table S12), and in contrast allowed for localisation of more relocations due to node parameters having lower horizontal asymptotes. Frequently, signals were detected but were not high enough to estimate location (Figure 6, Figure S11). Gaps between signal detections indicate that hummingbirds departed and returned to the grid’s detection area during the tracking period. Maximum duration of total time with tracking data was 97 days, while the time with signal detections stretched to 100 days for one male Great Sapphirewing that spent little time close enough to the grid to estimate continuous relocations (Figure 6).

Hummingbird tracks had a median speed of 0.08 m/s, with 95% of calculated speeds under 25 m/s. Any speeds greater than this were considered outliers and filtered out. Remaining points were below 18 m/s (95% CI), which we consider is a reasonable estimate of maximum horizontal velocity for these species although we do not know of studies that have calculated their flight power curves and maximum velocities. The average sizes of 95% utilisation distributions were 0.86 (± 0.55) km^2^ for Bronze-tailed Thornbills and 1.08 (± 0.26) km^2^ for Great Sapphirewings (Figure 6). All core utilisation distribution areas (50% UD) were located in parts of the valley covered by forest vegetation.

## Discussion

Movement data with high temporal resolution for small animals such as hummingbirds can be generated using an ARTS grid, but the accuracy in estimating locations must be considered carefully. In this study we assessed multiple decisions in the data processing workflow to produce localisation estimates and tested for the key spatial features that affect an ARTS grid’s accuracy. Our work provides guidance for the analytical decisions that must be made to produce location estimates. It is also an example of how ARTS grids may be used in ecosystems with complex vegetation and topography, and gives insight into what kind of questions may be addressed with this technology given its spatial resolution.

Radio telemetry is a powerful tracking tool that has been continuously improved by manufacturers to deploy very light tags that can give information on location for time frequencies of just some seconds and during prolonged periods (Naef-Daenzer et al., 2005). However, the implementation of ARTS as grids to track animals has lagged behind technological development (Ward et al., 2013), and the analytical methods to translate radio signal strength data to coordinates that constitute animal tracks are still diffuse (Paxton et al., 2022) even though they substantially affect error in localisation. However, ARTS are a promising solution for tracking small animals. We were able to collect high amounts of data on relocations within the area of our ARTS grid for calibration trials (2,361 total relocations) and tracked hummingbirds every 5 or 61 s for up to 97 days (with a total of 121,264 relocations for 9 individuals). Yet we also encountered technical difficulties and an average median localisation error of 105 m, which, until it can be further reduced, will continue to limit the use of this technology for some ecological research questions.

The main challenge in deriving estimates of location from radio signals is that radio waves may be blocked or attenuated by objects in their path between emission and reception. Consequently, signal propagation and resulting strength in natural settings is expected to be affected by vegetation, topography, or any other object, as well as environmental conditions such as temperature (Whitehouse et al., 2007). This makes the RSS-distance decay curve noisy, for signal strength does not only vary with distance. Moreover, in some instances, signal strength may be lower than the asymptote of the decay curve (Fisher et al., 2021) or signals may be detected by less than three nodes, rendering localisation impossible and resulting in loss of data. Our results show that this will be the case particularly when studying ground-dwelling animals, most probably because when a tag is close to the ground, emitted signals will be more heavily blocked by vegetation in comparison to when tags are higher up or even above vegetation height. Ward and Raim (2011) also encountered this difference in detection when tracking crows with an ARTS grid: the birds were not detected while they were on the ground but only when they flew up some metres in the air.

In this study, we obtained the lowest error when tags were flying above the height of the vegetation and we selected maximum RSS values, smoothed signals and tracks, used general model parameters, and conducted trilateration with the three strongest signal nodes. Selecting maximum RSS rather than average RSS values is preferable in a setting with bushes or trees because it will compensate for the attenuation of signals caused by vegetation, even though we excluded outliers that result from signal bouncing or multipathing when calculating average RSS. We have not encountered other studies that smooth RSS values, even though this step decreases the standard error of residuals in RSS-distance decay models. State-space models or spline smooths use the implicit autocorrelation in the data to smooth outlying values (Newman et al., 2023), given that values at one time step are influenced by values at previous or future time steps. We found that smoothing both RSS values and tracks reduced error. Future studies could further explore how movement models inherent in animal trajectories can further reduce error by incorporating predictions of future and past time steps.

Contrary to what we expected, using general model parameters for the RSS-distance decay curve produced better location estimates than fitting models individually by node or tag. Yet the effect magnitude of increased error from fitting models by node compared to general parameters was small, and we suspect that larger sample sizes (ensuring each node has many signal reads with more calibration trials) would have allowed better estimations with node-specific parameters and the effect of increased error may have even been reversed. In fact, we preferred estimating locations for tracked hummingbirds with node-specific model parameters to increase the number of relocations with location estimates, given that lower horizontal asymptotes in the RSS-distance decay relationship of some nodes resulted in the exclusion of fewer relocations.

Non-linear least squares is a fast and user-friendly approach to optimise the solution of multilateration, but it has previously been noted that using information from all receiving nodes could bias estimates of location by including nodes that receive distorted signals (Paxton et al., 2022, Luo et al., 2022). Although Paxton et al. (2022) had reported substantial decrease in error with filtering out nodes based on thresholds of RSS or distance in a flat landscape with low vegetation, we found that selecting the nearest nodes is actually the method that most increases error and is even less preferable than using all nodes. This result shows that localisation estimates that filter radio signals based on distance are not suitable in a landscape with more complex vegetation structure. In addition, using k-means clustering to introduce information from additional nodes may decrease error as suggested by Luo et al. (2022), but using only the three nodes with the strongest RSS reads is probably enough, as also mentioned in Wallace et al. (2022).

The influence of spatial features on localisation error is due to the attenuation of radio signals by vegetation and topography, key factors that should be considered when implementing ARTS grids in natural landscapes. In our site in the high Andes, vegetation is fairly open but shrubs and trees of the paramo and forest can grow to about 2–3 m high. Also, the valley’s ground elevation varies between 3,162 and 3,391 m asl (229 m difference). Both factors certainly increase estimated error in localisation, and explain differences with lower estimates of error for sites with almost no vertical vegetation structure, placement of receiving nodes higher up above the ground, and reduced distance between receivers (such as lowest median reported errors of 4.3 m in Wallace et al., 2022 and 28 m Paxton et al., 2022). Other studies for ARTS grids in landscapes of open areas interspersed with high vegetation, such as woodland, report higher median errors than what we found here (250 m in Lenske & Nocera, 2018, 322 m (average) in Scardamaglia et al., 2022). Placing nodes closer together should reduce error (Paxton et al., 2022), but is a trade-off with the area that may be covered and is limited by the project budget and duration given that more receivers will increase costs, effort, and time necessary for installation. Similarly, fine-scale radio-mapping of grids for methods such as that of Wallace et al. (2022) is logistically difficult for large grids in natural landscapes, and signifies an additional expense for projects. Our estimates for error could possibly change if more calibration trials were to be carried out. Also, exploring data-intensive methods such as machine learning approaches is an exciting future research avenue to improve accuracy of ARTS grids. In addition, the failure of one node during calibration of our grid highlights how ARTS grids require continuous maintenance, which is also a challenge for projects in remote areas and with limited staff available.

The error estimates that we found suggest that this technology is not appropriate for ecological research questions that require very high spatial resolution, such as studying interactions between individuals or species-specific foraging preferences. This ARTS grid is suitable to study broader-scale questions such as home range size, association with vegetation covers, and seasonality of occurrence. However, we found that error is lower for radio signals received above ground-level, suggesting that tracking studies in complex landscapes will be more suitable for flying animals than ground-dwelling ones (e.g. rodents, amphibians, lizards). Consequently, the selection of study species will also affect the performance of this technology.

In our pilot tracking of two hummingbird species, we estimated areas of utilisation distributions and recorded hummingbirds leaving and returning to the area. To the best of our knowledge, these species have never been tracked before and thus had no available movement data to make these kinds of inferences. In general, telemetry data for hummingbirds is scant because they are very difficult to track (but see examples of hand-tracking in Hadley & Betts, 2009, Hazlehurst & Karubian, 2018, Zenzal Jr. et al., 2018, among others), and thus very little is known about their movement ecology compared to other birds. Another study with high altitude Andean hummingbirds reports a similar territory size from handheld radio telemetry data (90th KDE average territory size for the Shining Sunbeam [*Aglaeactis cupripennis*] is 1.56 km^2^ [Pavan et al., 2020], vs. 95th AKDE of 1.08 km^2^ Great Sapphirewing and 0.86 km^2^ Bronze-tailed Thornbill reported here). Hand-tracking individuals is limited to short time periods (some days) and small areas, is strongly biassed towards areas that are easier for researchers to traverse (Hazlehurst & Karubian, 2018), and may not produce sufficient data points to estimate home range sizes (Ward et al., 2013). However, using an ARTS grid limits the spatial extent of localisation to the area where at least three receiving nodes from the grid detect a signal, therefore underestimating areas of use for animals that move out and away from the grid. Although we estimated localisations in the grid’s surroundings, radio signals detected by less than three nodes indicate birds were in the area but probably at a certain distance from the grid, whereas intervals with no radio signal suggest birds were even further away (Figure 6D-E).

The great advantage of ARTS grids is the very high amounts of data points that are produced over a longer period of time, giving additional insights into seasonal variation in movement. Although hummingbirds in the Andes have marked changes in seasonal abundance and altitudinal movement is likely widespread (Barçante et al., 2017), so far these patterns have been documented by comparing changes in abundance through monitoring different elevation bands through time (e.g. Gutiérrez Z. et al., 2004) or using citizen science data to estimate within-year changes in species’ distributions (Rueda-Uribe et al., 2024). We anticipate that through continued tracking of hummingbirds with the ARTS grid, we will collect more information on their seasonal occurrence in this site (as in Smetzer et al., 2021), since the available food resources of high Andean ecosystems are variable and respond to flowering pulses mainly mediated by rainfall (Pelayo et al., 2021). Core areas of use (50% UDs) were all located in parts of the valley were elfin forest shelters plants that are visited by the two species we tracked and were in flower between December and April (e.g. *Bomarea*, *Bejaria*, *Macleania,* and *Tibouchina*). These patterns may change when the plants of the paramo and open grassland areas (*Espeletia* and *Puya*) flower later in the year, and can be quantified with habitat-selection analyses (Fieberg et al., 2021).

## Conclusion

Our study presents an assessment of how the accuracy of ARTS grid is affected by analytical decisions while processing radio signals for localisation as well as the spatial features of a natural landscape. We tested this technology by tracking challenging subjects, thus providing proof of concept for investigating movement across multiple habitats of small animals; successfully studying two species of high altitude hummingbirds that previously lacked movement ecology data. It therefore provides guidelines for researchers setting up other ARTS grids to study wild animals, reiterating the importance of extensive calibration before tag deployment but also highlighting the variety of decisions that may be undertaken during the data workflow. The comprehensive evaluation of the error produced by different approaches also gives insight into what kind of research questions may be currently approached with this technology, particularly in sites with vertical vegetation structure and rugged terrain. One of the challenges we encountered while working in a remote area is that ARTS grids require continuous maintenance and supervision, and so tasks such as fixing failing equipment or cleaning the solar panels that provide receiving nodes and antenna with energy become difficult. Our approach could possibly be improved in the future by increased calibration trials for data-intensive approaches with machine learning. The spatial structure of radio signals and the movement properties of animal tracks could be modelled to reduce error, although this is a trade-off with higher project costs and time.

## Acknowledgements

We are grateful to all students, park rangers, and volunteers that supported and participated in fieldwork. They braved through the extreme weather of the paramo and a muddy valley with challenging slopes and prickly vegetation, carrying heavy equipment and doing intense physical work. We are particularly thankful to Juan Pablo Ríos, Manuela Lozano, Jonathan Espitia, Sarah Chaves, Angie Rodríguez, Ana Melisa Fernandes, Miguel Ángel Muñoz-Amaya, Aaron Skinner, Daniel Mancera, Fredy García, Luisa Díaz, Ángela María Uribe, Alfonso Rueda, Adriana Rueda, Diego Rueda, Michael Spence and Santiago Cepeda. We also thank our drone pilots, Nicolás Skillings and Carolina Arévalo, Jessie Williamson for sharing experiences in hummingbird tracking and research, Luis Guillermo Linares for giving initial advice and support, Leonel Herrera-Alsina for help running code on the cluster, Stephen Palmer, Rebekka Allgayer and Tamsin Woodman for suggestions on the methods, David Sragli, Rufus Behr and Alice Scarpa for initial coding approaches, Chingaza National Natural Park staff and management, especially Juan Camilo Bonilla-C., for all the support in the field, and Premium 3D for designing a tag chassis specifically suited for hummingbirds. NERC, Rufford Foundation, University of Washington Department of Biology, and School of Biological Sciences University of Aberdeen funded this work and we are grateful for their support.

## Funding

This work was supported by the UKRI Natural Environment Research Council (grant no. NE/S007377/1 to C.R.U.), the Rufford Foundation (Small Grant no. 36476-1 to C.R.U.), University of Aberdeen School of Biological Sciences (Charles Sutherland Scholarship Fund, to C.R.U., and Windfall grant to J.M.J.T), University of Washington Department of Biology endowed funds (Orians Award for Tropical Studies, Margo & Tom Wyckoff Award, to A.J.S.), and the Walt Halperin Endowed Professorship and the Washington Research Foundation as Distinguished Investigator (to A.R-G).

## Conflicts of interests

The authors report no conflict of interest.

## Data availability

All data and code will be made available when this manuscript is peer-reviewed.

## Statement on inclusion

Our study included a group of people with a diversity of disciplinary backgrounds and career stages during project design, fieldwork, analyses and writing. People involved are from different countries, and especially from or based in the country where fieldwork was carried out.

## Notes

### Competing Interest Statement

The authors have declared no competing interest.

