## Supplementary methods and figures for "Tracking small animals in complex landscapes: a comparison of localisation workflows for automated radio telemetry systems"

*Grouped multilateration with k-means clustering*

Grouped multilateration with k-means clustering consists of producing multiple location estimates per relocation by using all the possible combinations of subgroups of nodes that detected a signal (where group sizes must be of 3 nodes or more). All possible location estimations from groups of at least 3 nodes of a total of n nodes is given by the equation

$$\sum_{r=3}^{n} C(n,r) \;=\; \frac{n!}{r!(n-r)!}$$

where r is the number of nodes that may be used to estimate each location and n is the total number of nodes that received a signal by relocation. This means that a relocation with only 3 nodes will only have one estimated location, while a relocation with 40 nodes will have 1.099512e12 estimated locations. To reduce computing time, we set a threshold of a maximum of 7 nodes to be used in each estimation, meaning that a maximum of 99 possible estimates by relocation were used for k-means clustering. This threshold was set by inspecting how the maximum RSS detected by a top node decreased with consecutive nodes that detected a signal (Figure SM1). We found that on average after the 5th node, reduction in RSS reaches an asymptote, meaning that after the 5th node additional nodes are possibly less informative.

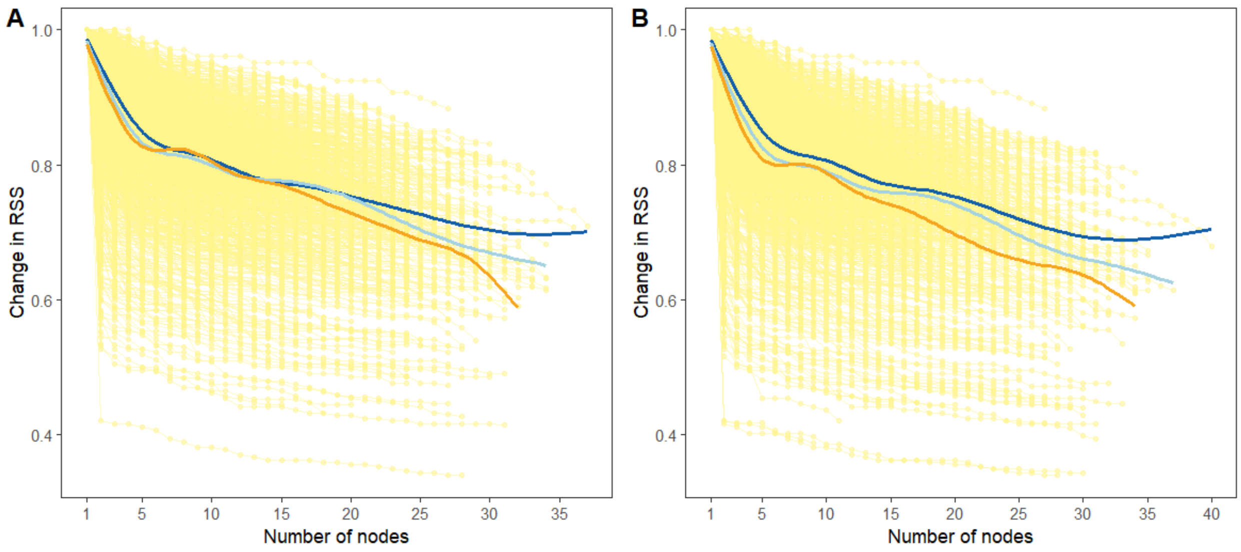


**Figure SM1**. Change in radio signal strength (RSS) for each relocation considering additional nodes after the strongest signal node. The first strongest signal node will have a value of 1.

Points indicate individual nodes and relocations are connected with lines. Thicker lines show trend with a loess smoother of calibration trials at high flights (dark blue), low flights (light blue) and ground tests (orange). Panel A) shows values for average RSS and B) for maximum RSS.

The selection of an optimal value for k is based on the reduction of total within cluster sum of squares (wss), choosing the smallest value of k within the 95% confidence interval of the estimated asymptote (Figure SM2). K-means clustering was done with Hartigan-Wong algorithm (Hartigan and Wong, 1979), using 10 random starts. If the number of nodes that detected a signal was 3, only one location may be estimated (there is only one possible group with minimum 3 nodes) and therefore we did not proceed with k-means clustering but instead kept the only possible location estimate. In the case that only 4 nodes detect a signal, there are only 5 possible location estimates, and therefore calculating the asymptote of the decay relationship between wss and k is unrealistic. In these cases, we set k as 3.

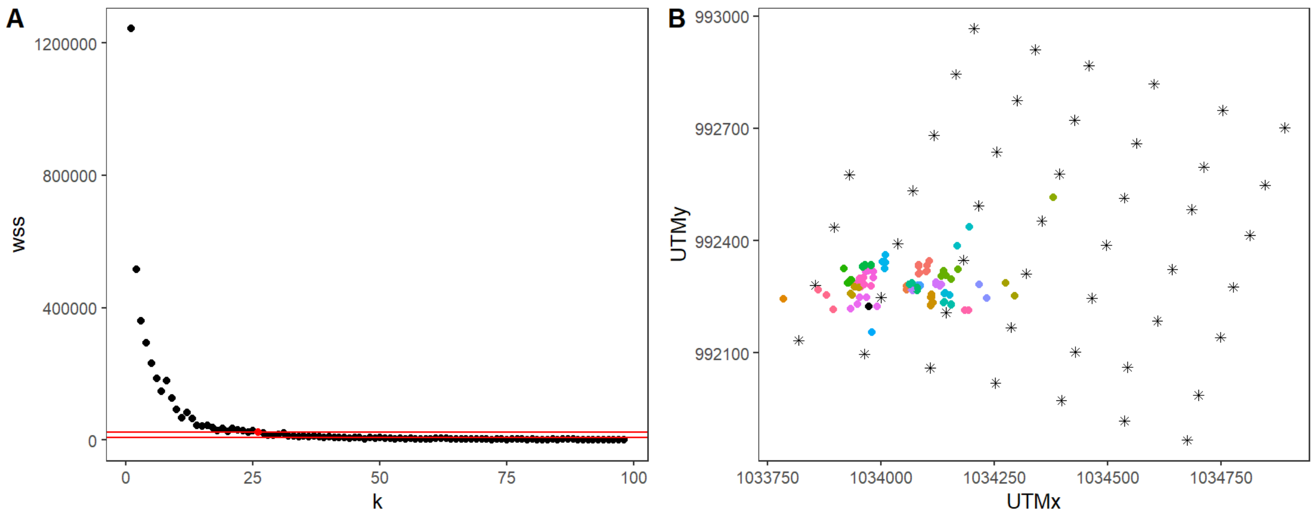


**Figure SM2.** A) Exponential decay of total within cluster sum of squares (wss) with varying number of clusters (k). Red lines show 95% confidence interval of calculated horizontal asymptote, and red dot indicates chosen optimal k (k=26) for this example relocation. B) True location of this example relocation is shown by the black point within the grid, where nodes are represented by asterisks and are approximately 150 m apart. This relocation was detected by 7 nodes and therefore has 99 possible location estimates, which are shown as coloured points. Clustering with k=26 is indicated by the colours.

***Factors that influence error : linear mixed effect models - LMMs***

To compare the analytical and spatial factors that influence error, we fit linear mixed effects models (LMM).

In the case of the LMM to assess how analytical decisions influence error, we included calculated error as the response variable, each relocation nested within its calibration trial as random effects and all steps in data processing as fixed effects:

> *Error ~ Signal selection + Signal smoothing + Model parameters + NLS multilateration method + Track smoothing + (1 | calibration trial/relocation)*

Error was transformed by the square root to reach normality and the inclusion of random effects was sufficient to reduce temporal autocorrelation (Figure S7). We compared all the possible combinations of including fixed effects in the models (32 including null) and ranked resulting models with the corrected Akaike Information Criterion (AICc).

For the spatial LMMs, the response variable was calculated error, calibration trial was the random effect and the fixed effects were the spatial features of ground elevation, terrain ruggedness, distance to the edge of the grid, presence inside or outside the grid, flight height and type of vegetation cover, as well as all the interactions between the three vegetation cover types and the first four landscape variables. Since ground elevation and terrain ruggedness were correlated (Figure S8), from these two we only included terrain ruggedness as the more informative variable in the models. The final fitted full model included the following terms:

> *Error ~ terrain ruggedness + distance from edge + inside or outside grid + flight height + dominant vegetation cover + terrain ruggedness*dominant vegetation cover + distance from edge*dominant vegetation cover + inside or outside grid*dominant vegetation cover + flight height*dominant vegetation cover + (1 | calibration trial)*

In this model, error was also transformed by the square root to reach normality and outliers (under 2.5%, 15 m and over 97.5%, 410 ) were removed. All quantitative fixed effects (terrain ruggedness, distance from edge, flight height and proportion of vegetation covers) were scaled around their mean and transformed by the square root. The inclusion of calibration trial as random effect was not sufficient to reduce temporal autocorrelation in normalised residuals (Figure S9), so before fitting the final model we thinned the data by removing every two time lags. Using the final thinned model, we assessed all the possible combinations of fixed effects in the models (97 models including null) and ranked models according to AICc. Estimated coefficients and confidence intervals were calculated by the full averaging of highest-ranking models with AICc weights (top 3 models, summed 77% AICc weights and AICc delta from 0.99 to 2.93 to fourth highest-ranking model).

### *Continuous-time movement models: track simulation and utilisation distribution estimates*

Continuous-time movement models (CTMMs) were fitted by calibration track or individual tracked animals. Top-ranking models were selected using “ctmm.select” function in the “ctmm” package in R (Calabrese et al., 2016, Fleming & Calabrese, 2023). This function fits different movement models that consider autocorrelation in position and velocity as well as limits in range residencies and either isotropic or anisotropic directions of diffusion (Fleming & Calabrese, 2023). Briefly, the independent identically distributed model (IID) assumes no autocorrelation, Brownian motion models diffusion with no autocorrelation in velocity nor home range limitations, and Ornstein-Uhlenbeck processes can incorporate range residencies (OU), autocorrelation in velocity (integrated - IOU) or both (foraging-OUF) (see Calabrese et al., 2016).

We used fitted CTMMs of calibration tracks to simulate tracks within the automated radio telemetry grid to compare estimated locations to those expected from a random process. For this we fitted CTMM models, incorporating a known maximum error of 5 m (recorded by handheld GPS, for drone flights the manufacturer does not report accuracy but it is assumed to be within the same range or lower), and setting the number of iterations to 99 for each track. Resulting simulations that were located outside of the grid plus a 100 m buffer were excluded, and simulated tracks within the grid were allowed to rotate in any direction and randomly be displaced anywhere within 100 m of the area covered by the grid. Differences between estimated random locations and known locations were calculated as the distance in metres.

In addition, we fitted CTMMs to recorded hummingbird tracks to calculate kernel density utilisation distribution areas that account for autocorrelation (Silva et al., 2022). For these fitted models, we set the error to the median calculated error of flying trials (both high and low flight, 97 m).

**Supplementary figures**

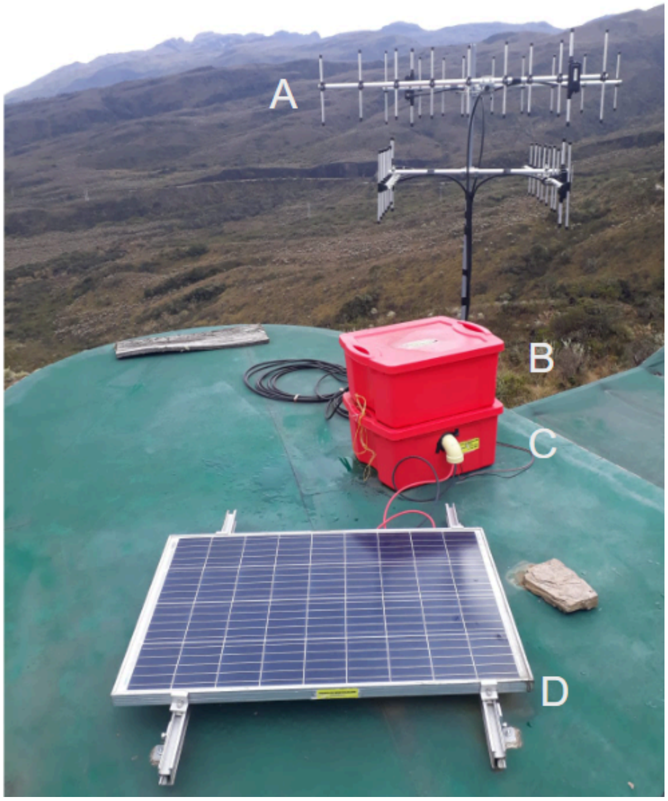


**Figure S1**. Antenna setup located approximately 10 m above the ground on the roof of a park rangers' cabin overlooking the Valle de los Frailejones. A) 4 Yagi antennas, B) box that houses the sensor station, C) box that houses deep cycle marine battery and D) solar panel to replenish the battery.

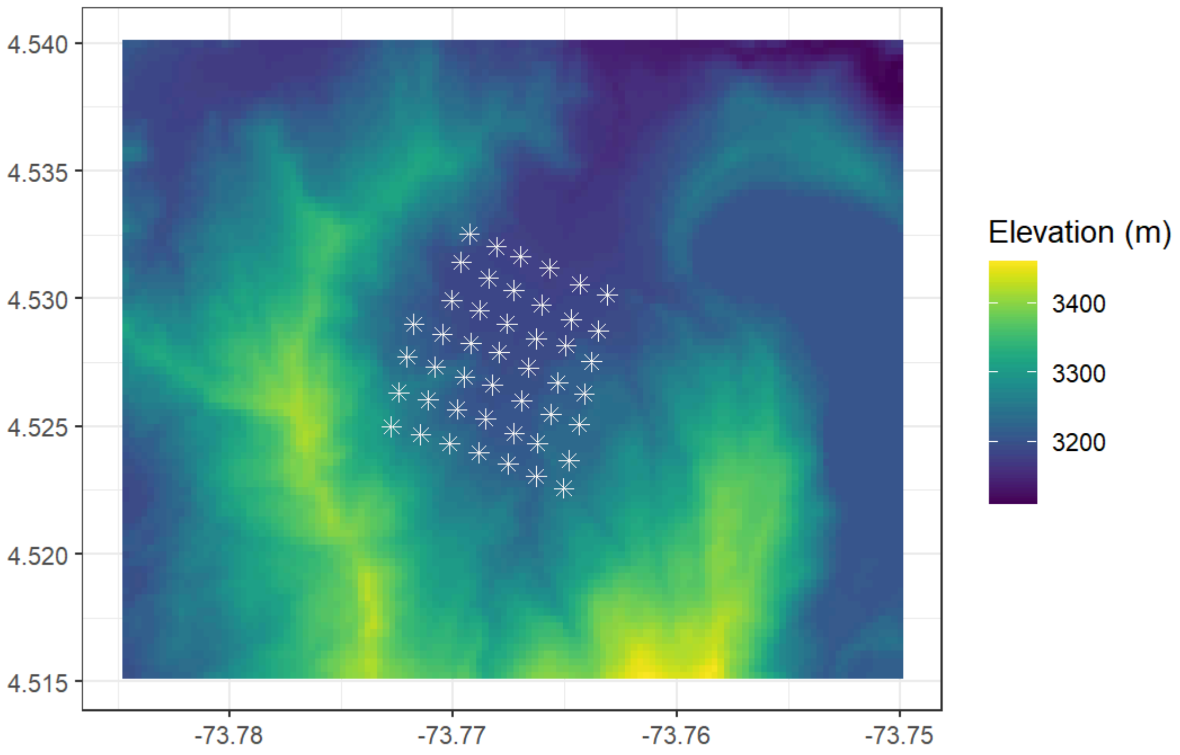


**Figure S2**. Ground elevation in area of automated radio telemetry grid in the Valle de los Frailejones in Chingaza National Natural Park. The 46 receiving nodes installed in the valley are shown as white asterisks, and are set approximately 150 m apart. Data was downloaded from GLO-30 Copernicus digital elevation model (30 m resolution) of the European Space Agency (ESA, 2024).

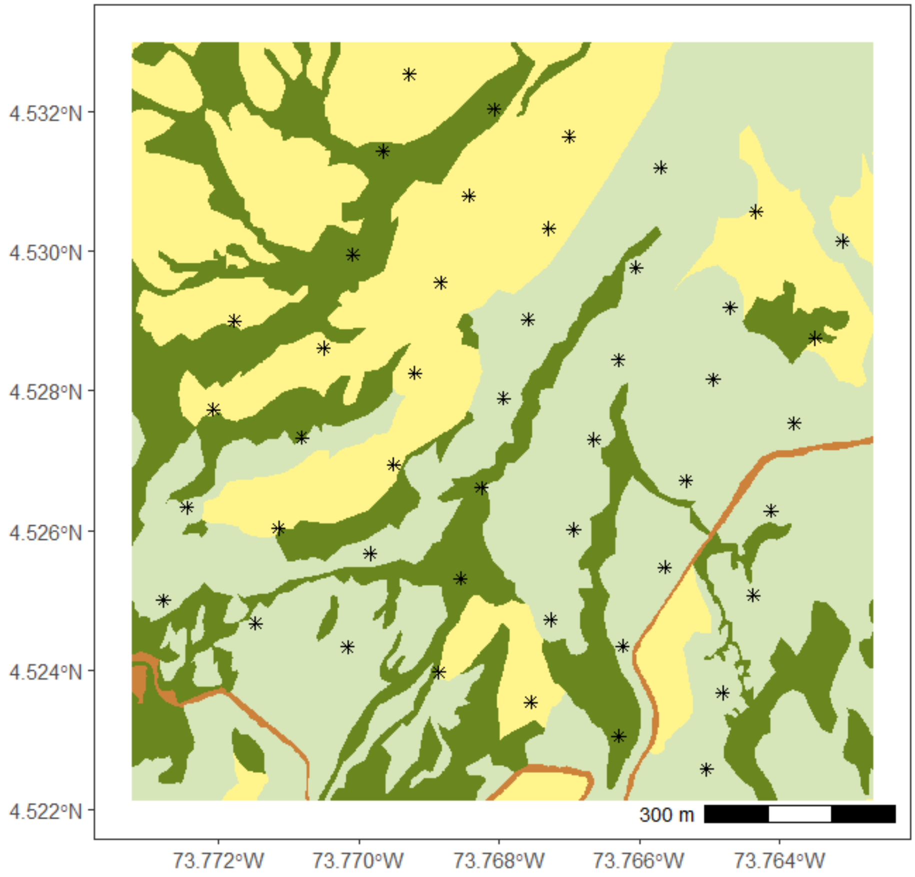


**Figure S3**. Vegetation classification map of the automated radio telemetry (ARTS) grid, at 1 x 1 m cell resolution for area within 50 m of grid. Dark green shows more dense vegetation of forest and bamboo, light green paramo, yellow open grassland and orange built-up areas (dirt road and park ranger cabins). The 46 receiving nodes that are set up are shown as asterisks.

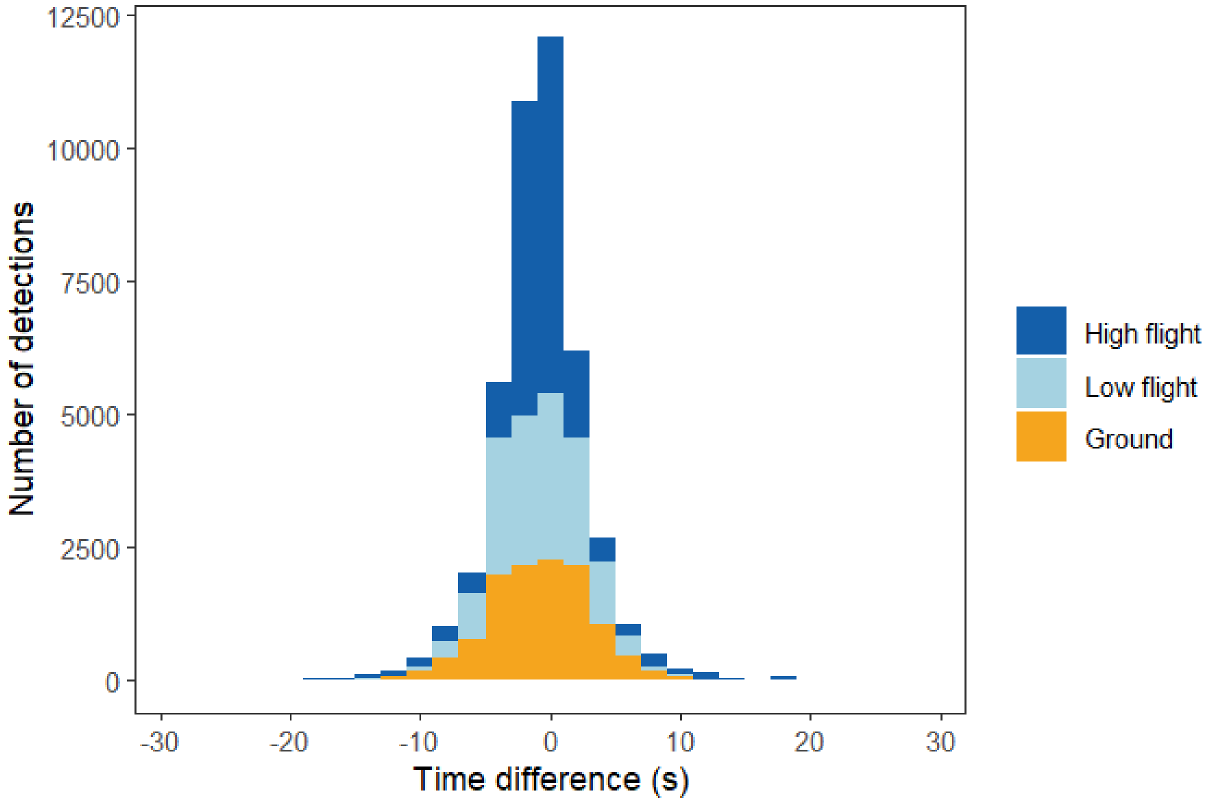


**Figure S4.** Time difference in seconds between tag relocation and signal detection during calibration trials at high flight (dark blue), low flight (light blue) and ground level (orange).

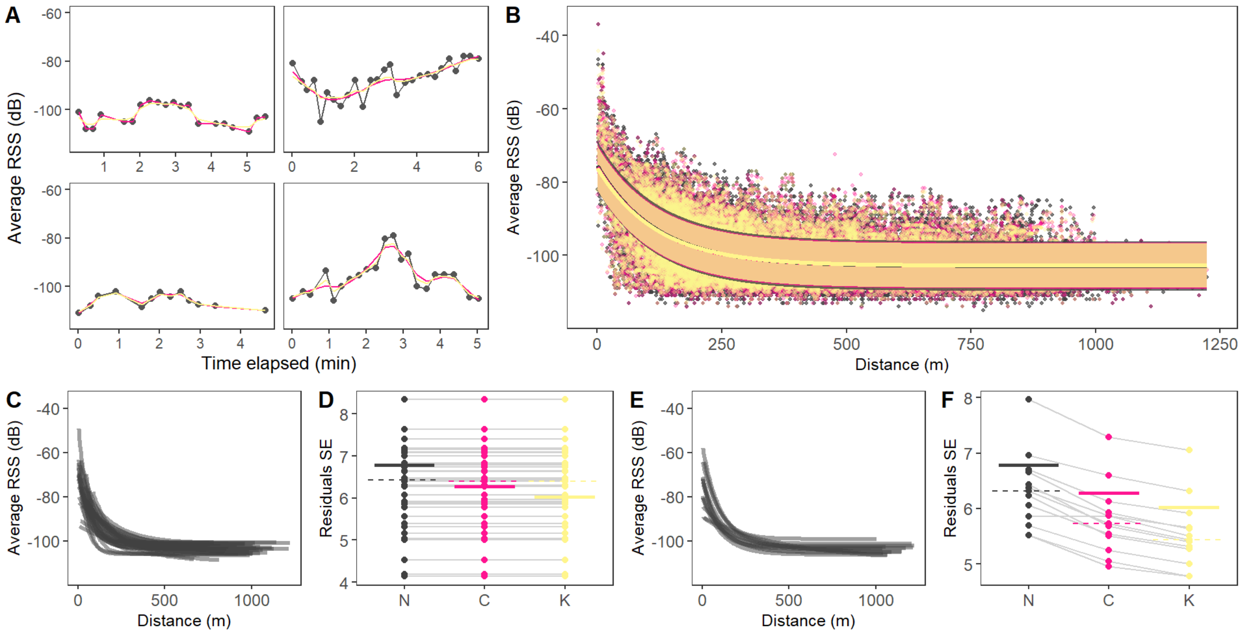


**Figure S5**. Detected average radio signal strength (RSS) and variation of fitted exponential decay models. Colours throughout all panels represent average RSS values that were not smoothed (black), smoothed with cubic splines (pink), or Kalman smoothers (yellow). A) Example of average RSS detected for one calibration trial by four different receiving nodes through time. Points show initial reads and lines smoothed values with cubic splines or Kalman smoothers. B) Relationship between average RSS and measured distance for calibration trials. Line shows fitted non-linear least squares exponential decay models and shaded area indicates standard error (SE) of residuals. Models fitted individually by receiving C) nodes or E) tags, each line represents a separate node or tag. Standard error of residuals in models by D) node or F) tag, for maximum RSS values that were not smoothed (N), smoothed with cubic splines (C), or Kalman smoothers (K). Points show individual receiving nodes or tags and lines connect the same node or tag. Solid thick horizontal lines show SE for general model and dashed horizontal lines show median value for models separated by nodes or tags. See Figure 4 for graphs of maximum RSS.

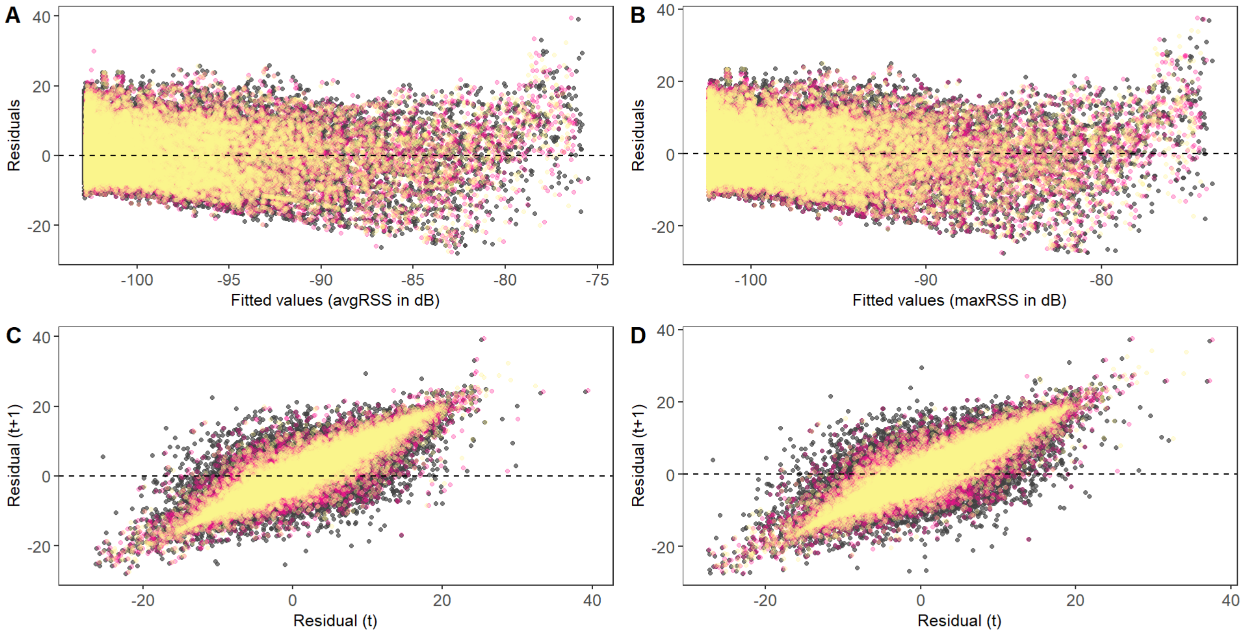


**Figure S6**. Scatterplots of normalised residuals and fitted A) average and maximum B) radio signal strength (RSS) for non linear least square (NLS) exponential decay models. Lower panels show normalised residuals at time t and t+1 (next time step) for models of C) average RSS and D) maximum RSS. Colours indicate when signals were not smoothed (black) or smoothed by cubic splines (pink) or Kalman smoothers (yellow).

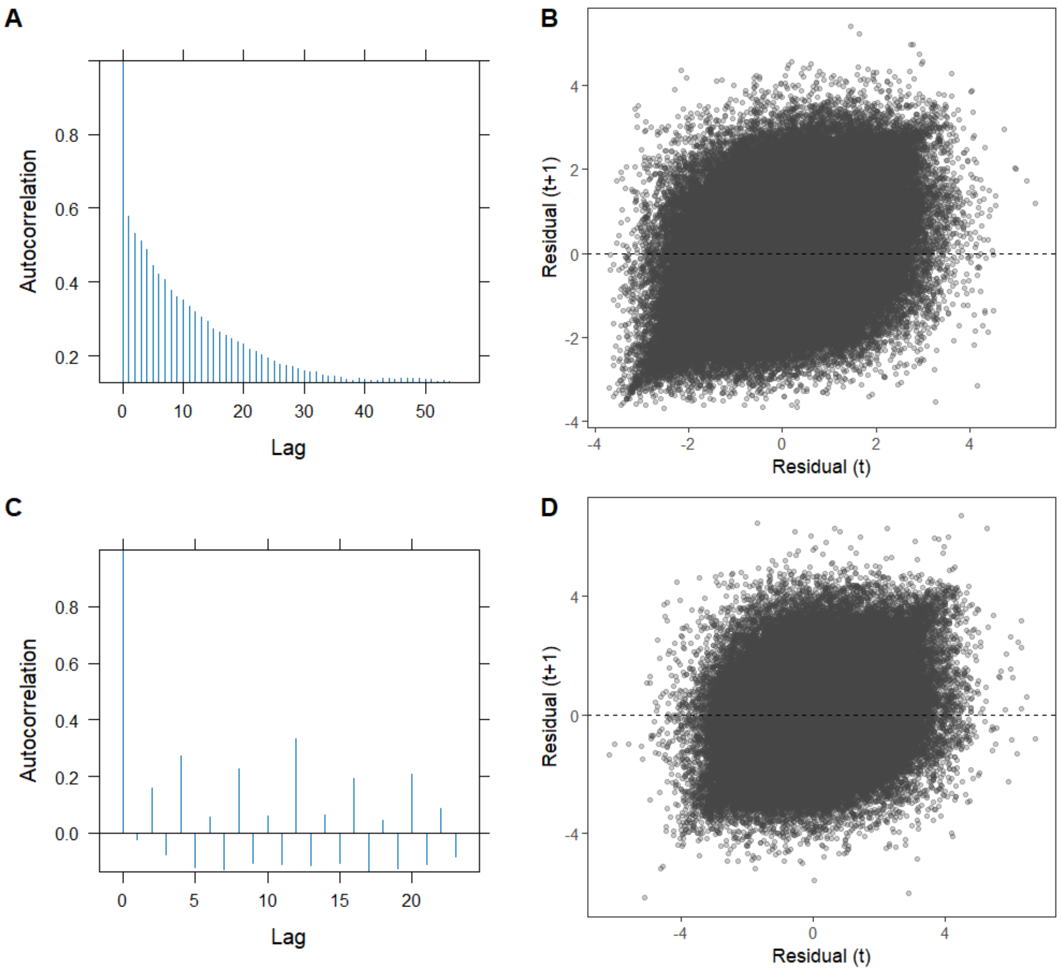


**Figure S7**. Autocorrelation correlograms and scatterplots of normalised residuals at time t and t+1 (next time step). Panels A and B show residuals of models when relocations within trials were not included as random effects, and panels C and D when relocations within trials were included as random effects.

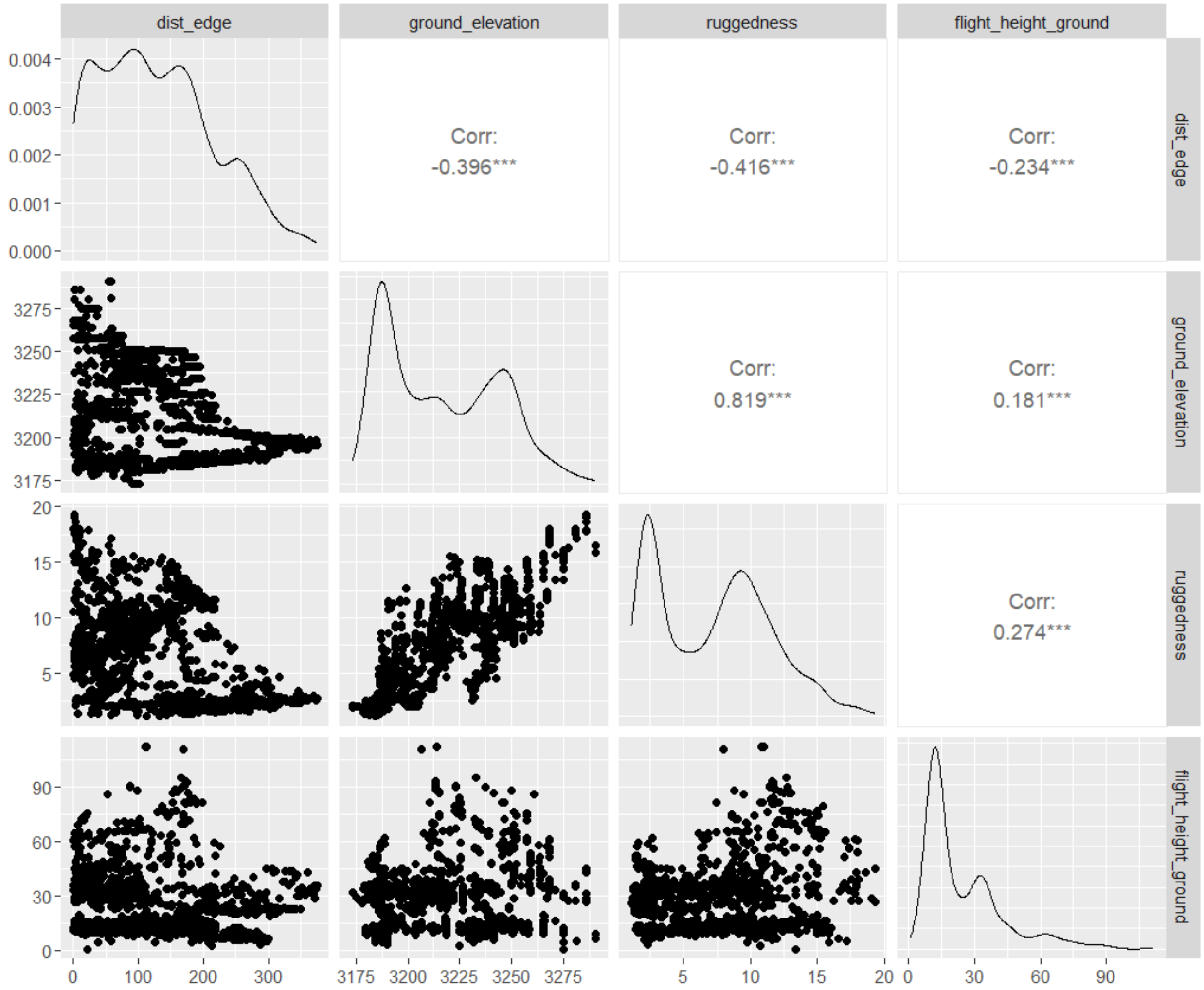


**Figure S8.** Correlation between fixed effects included in linear mixed models for the influence of spatial features in localisation error. dist_edge = distance from the edge of the grid. Ground elevation removed because it is highly correlated with terrain ruggedness.

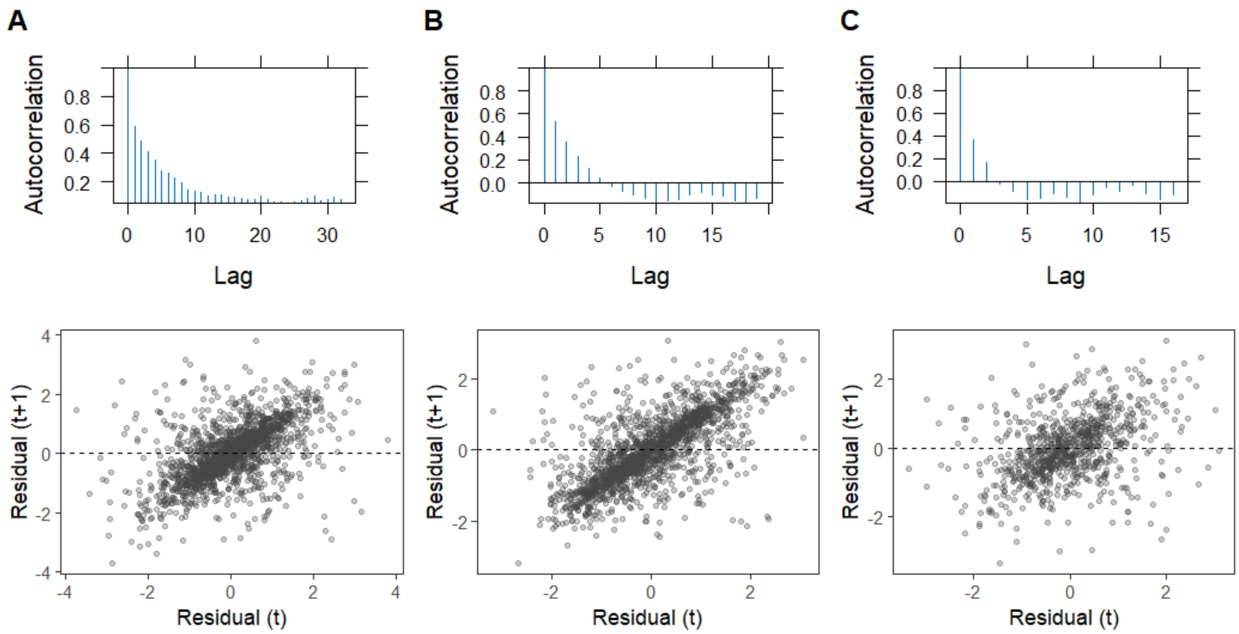


**Figure S9.** Autocorrelation correlograms and scatterplots of normalised residuals at time t and t+1 (next time step). Panel A shows residuals of models when trials were not included as random effects, panel B whentrials were included as random effects and panel C when data was thinned by removing two observations between neighbouring relocations.

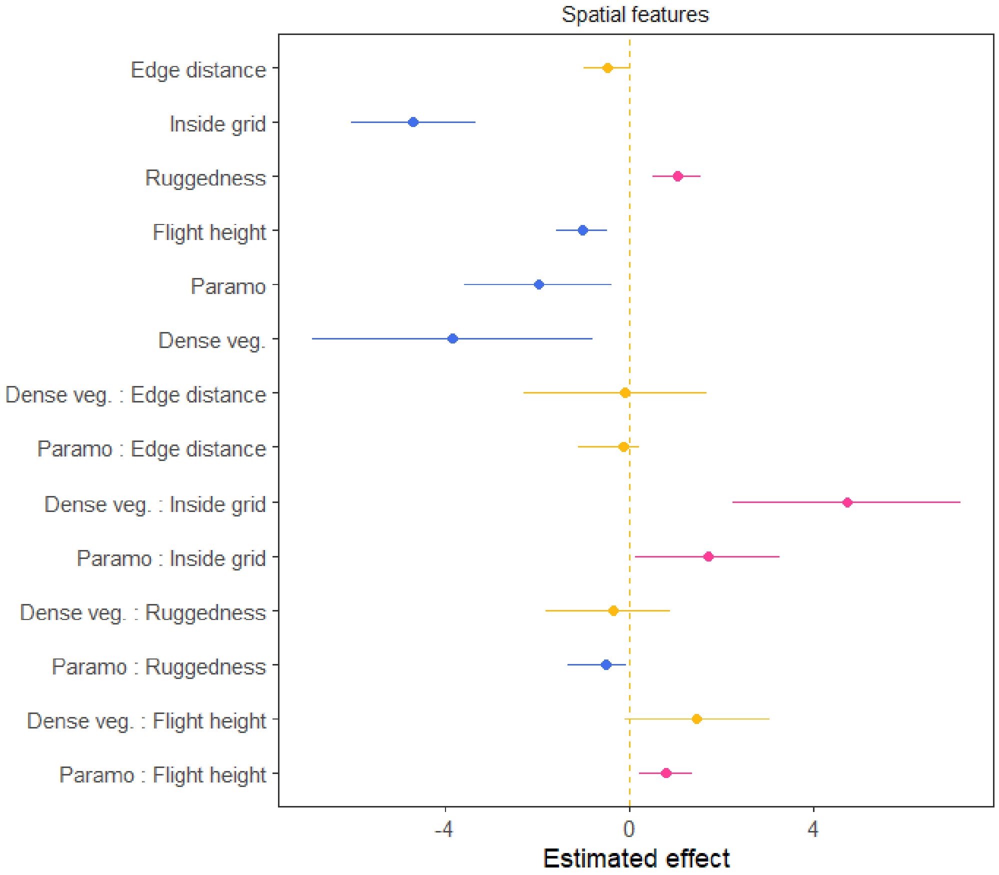


**Figure S10.** Estimated effects for linear mixed models of the influence of spatial features on localisation error. Points are estimated effects and horizontal lines 95% confidence intervals (CI), with the vertical dashed line showing an effect of zero. Yellow indicates estimates with 95% CI that overlap with zero, while blue and pink show negative effects (reduced error) or positive effects (increased error), respectively. Effect magnitude shown has the square root transformation to keep model assumptions of normality. Two points between vegetation covers and first four spatial features indicate interactions. Note that the vegetation cover of open grasslands was chosen as baseline for comparison in the models.

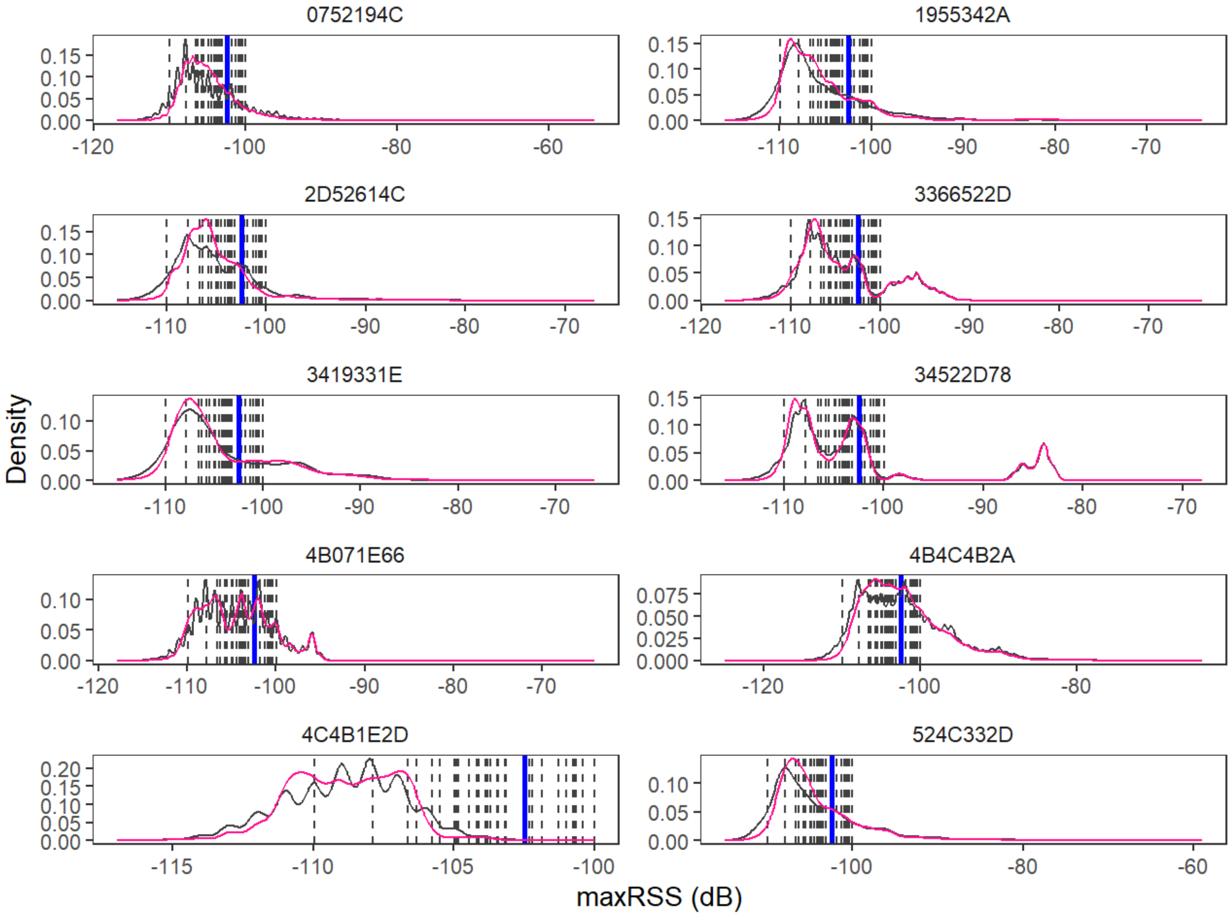


**Figure S11.** Distribution of recorded maximum radio signal strengths for each individual tagged hummingbird. Values without smoothing are shown by the black curve and values that were smoothed with cubic splines are in pink. Vertical lines show horizontal asymptotes for estimated decay function relationship between recorded RSS and distance, thus showing the minimum RSS value that may be used to estimate distance of a relocation from a node. Dashed black lines show asymptotes calculated for nodes, solid blue lines for models with general parameters (-102.47 dB).

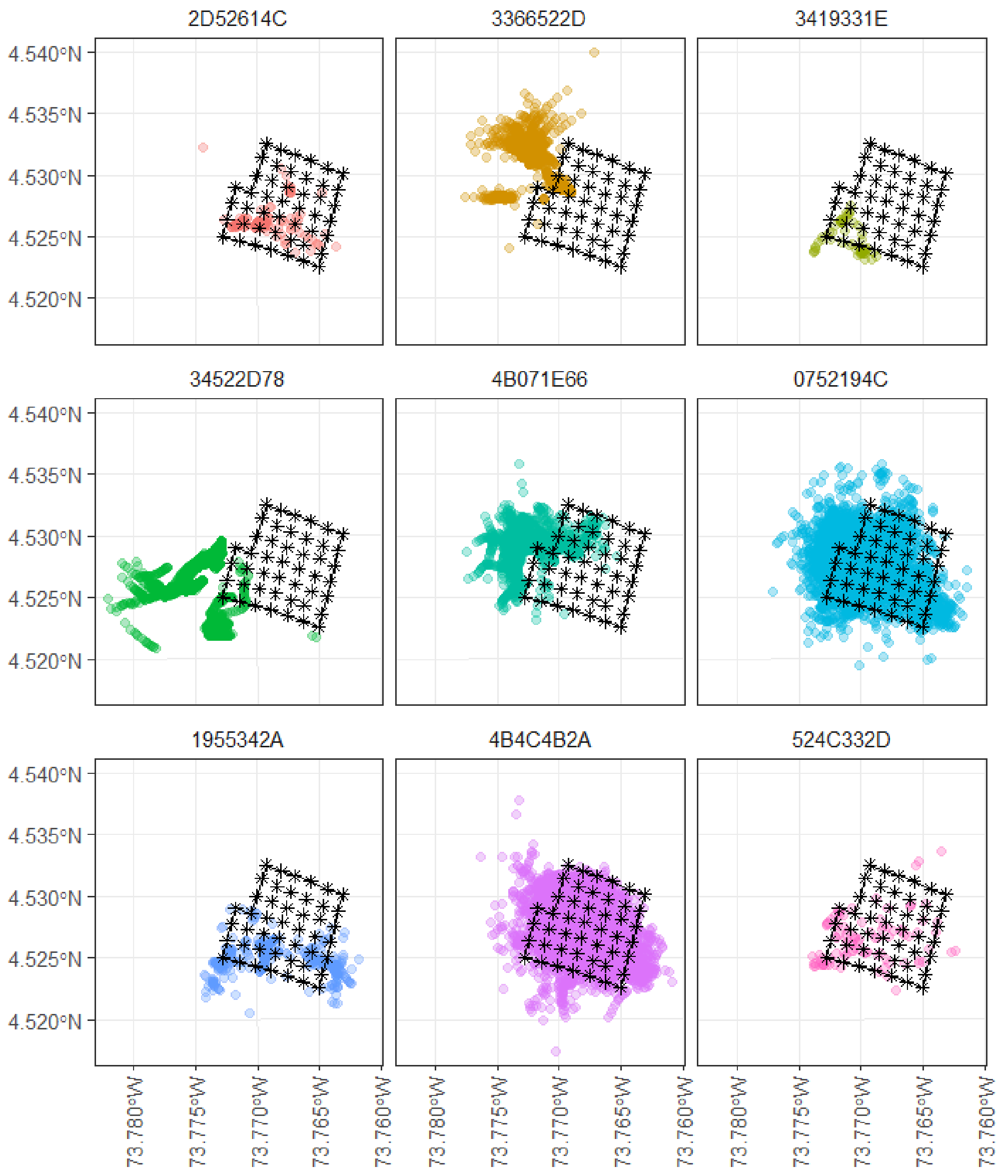


**Figure S12.** Estimated localisations of 9 tracked hummingbirds. Asterisks indicate receiving nodes.

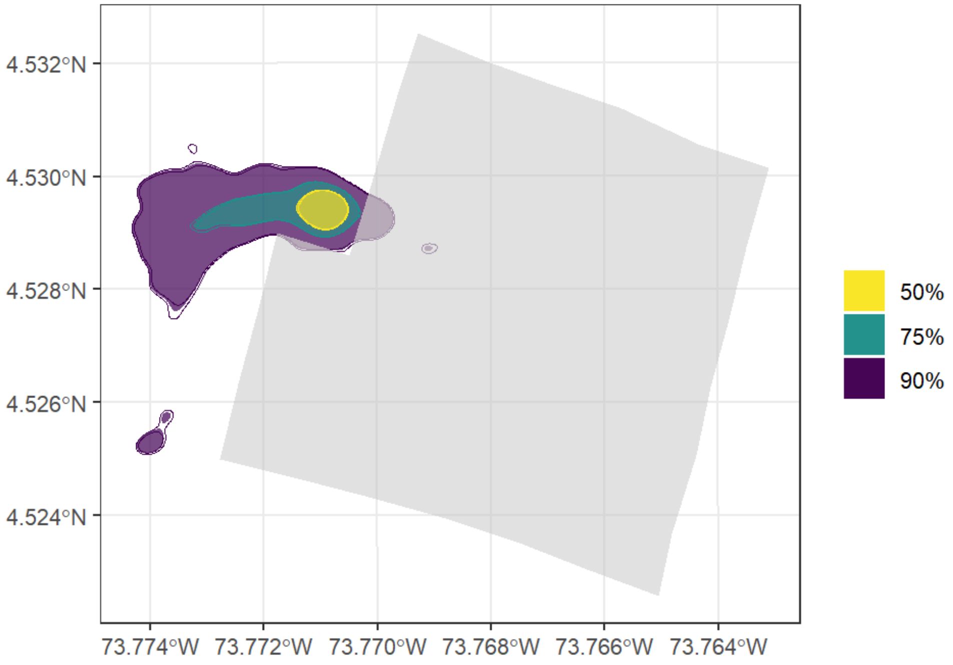


**Figure S13.** Utilisation distribution measured with autocorrelated kernel density estimates (AKDEs) of one Bronze-tailed thornbill that was much smaller than the rest. Filled polygons show estimates for 50, 75 and 95% AKDEs, with lines indicating lower (inner line) and upper (outer line) 95 % CI for each. Light grey polygon indicates the area covered by the automated radio telemetry grid.
